# A metagenomic-based study of two sites from the Barbadian reef system

**DOI:** 10.1101/2021.01.02.425083

**Authors:** S Simpson, V Bettauer, A Ramachandran, S Kramer, S Mahon, M Medina, Y Valles, V Dumeaux, H Valles, D Walsh, MT Hallett

## Abstract

We study the microbiome of sea water collected from two locations of the Barbadian coral reefs. The two sites differ in several environmental and ecological variables including their endogenous benthic community in addition to their proximity to urban development and runoffs from inland watersheds. The composition of the microbial community was estimated using whole genome DNA shotgun sequencing. Although both sites exhibit a similar degree of richness, the less urbanized site (Maycocks reef at Hangman’s Bay) is strongly concentrated with phototrophs. In comparison, the more urbanized location (Bellairs Research Institute) is enriched for copiotrophs, macroalgal symbionts and marine-related disease-bearing organisms from taxa scattered across the tree of life. Overall, our samples and associated measurements of chemical and environmental qualities of the water are in line with previous marine microbiome profiles of warm ocean surface waters. This suggests our approach captures salient information regarding the state of each coral reef site and sets the stage for larger longitudinal studies of coral reef dynamics in Barbados.

## Introduction

Coral reefs are fragile ecosystems which house a significant fraction of the planet’s biodiversity and serve as the economic foundation to many communities, providing a source of food and a defense mechanism against extreme meteorological events.^1-5^. Tropical coral reef systems require precisely tuned conditions for efficient ecological functionality and survival including clear oligotrophic water^6,7^, a narrow band of water temperatures^8^, larval recruitment from distal origins^9^ and continuous nutrient uptake^10^. Tropical coral reef systems in the Caribbean region have been subject to both global and local stressors over the past 50 years, resulting in an 80% loss of coral cover^11,12^. Global stressors include climate change with associated increased sea temperatures and severe storms^1,3,13^, ocean pollutants including plastic waste^14^ and ocean acidification^15^.

The reef system studied here, which lies on the west side of Barbados, has witnessed a 70% decline in coral colonies over the past two decades and has suffered two mass bleaching events in 2005 and 2010^11,16-18^. Barbados has also experienced repeated golden tidal events caused by Sargassum seaweed originating from the tropical Atlantic ocean, east of Brazil, which stress the ecosystems via light reduction, oxygen depletion, and by increasing nutrient levels and mortality of associated reef fishes^19,20^. These same reef systems are negatively impacted by several biotic and abiotic local stressors including overfishing^12^, mechanical disturbances from water-based activities, and eutrophication due to agricultural runoffs, sewage, and anthropogenic pollution^11,21,22^

Since microbiomes influence and reflect the environment they inhabit, an understanding of the natural variability and shifts in community gradients of the coral reef sites may allow for the precise identification of environmental disturbances, in turn suggesting specific prophylactic measures that could be taken to cull negative influences^23-27^. Indices of reef health have been traditionally formed by combining information obtained from physical ecological assessments (tissue damage, coral disease and bleaching)^28,29^ and measures of diversity of fish and other reef inhabiting species^30,31^. The status of the Barbados reef system has relied on such methods since 1982 to monitor 43 sites in five-year cycles^17^. The microbial community within the corals and water surrounding these corals has not been studied extensively nor been subjected to high-throughput unbiased molecular profiling. Our goal here is to profile the Barbadian coral reef microbiome and integrate this information within the context of similar efforts led by international consortia to globally profile oceanic microbial communities^8,32-34^.

These efforts exploit the power of high throughput metagenomics sequencing in order to characterize coral reef systems, and show how features change due to biotic and abiotic influences, unravelling key links to environmental variables^27,35^. The characterization of the microbiomes across varying ecological, environmental and geophysical properties in Barbados represents a first step towards the identification of molecular biomarkers of reef health progression and adaptation, and also towards identifying interventions for coral reef restoration.

## Results

### The two reef sites differ in a range of ecological, environmental and chemical metrics

The Bellairs reef lies off the west (Caribbean) coast of Holetown, a densely populated area of the island with tourism and other urban infrastructure^36^. It is subject to several runoffs originating from an inland watershed^37^. The Bellairs reef lies within the Babados (Folkestone) Marine Reserve and is protected from fishing and vehicle traffic, although still subject to high recreational activity. The Maycocks reef also lies off the west coast but in a sparsely populated area with no coastal urban development but a small residential community inland^36^. Whereas the Bellairs site is the back reef zone of a fringing coral reef directly adjacent to the shore in shallow water (∼10 m), Maycocks is a bank reef located in deeper water (∼15m) approximately 1.5 km from shore (**Figure 1A-C**). The benthic composition at both sites was determined after both recent bleaching events but two years previous to this study^11^. In comparison with Bellairs, Maycocks exhibits a higher abundance of corals, a more uniform distribution of species, significantly more hard corals, less filamentous algae, a lower average percentage of dead corals and greater average fish biomass **(Table 1A)**. Water samples were collected from both sites ∼1m above each reef, and sequentially filtered to concentrate organisms between 3µm and 0.22µm in line with Tara Oceans parameters^33^. We measured several chemical and environmental parameters and observed that Maycocks had a significantly greater level of dissolved oxygen and lower level of nitrate (NO_3_; **Table 1B, Methods 1**,**2**). We also estimated the number and size of microbes present in our samples using microscopy (**Methods 2**) and observed that overall both sites had approximately the same number of organisms per mL^3^, but that Maycocks has a tendency towards smaller organisms compared to Bellairs (**Supplemental Figure 1**).

**Table 1.**
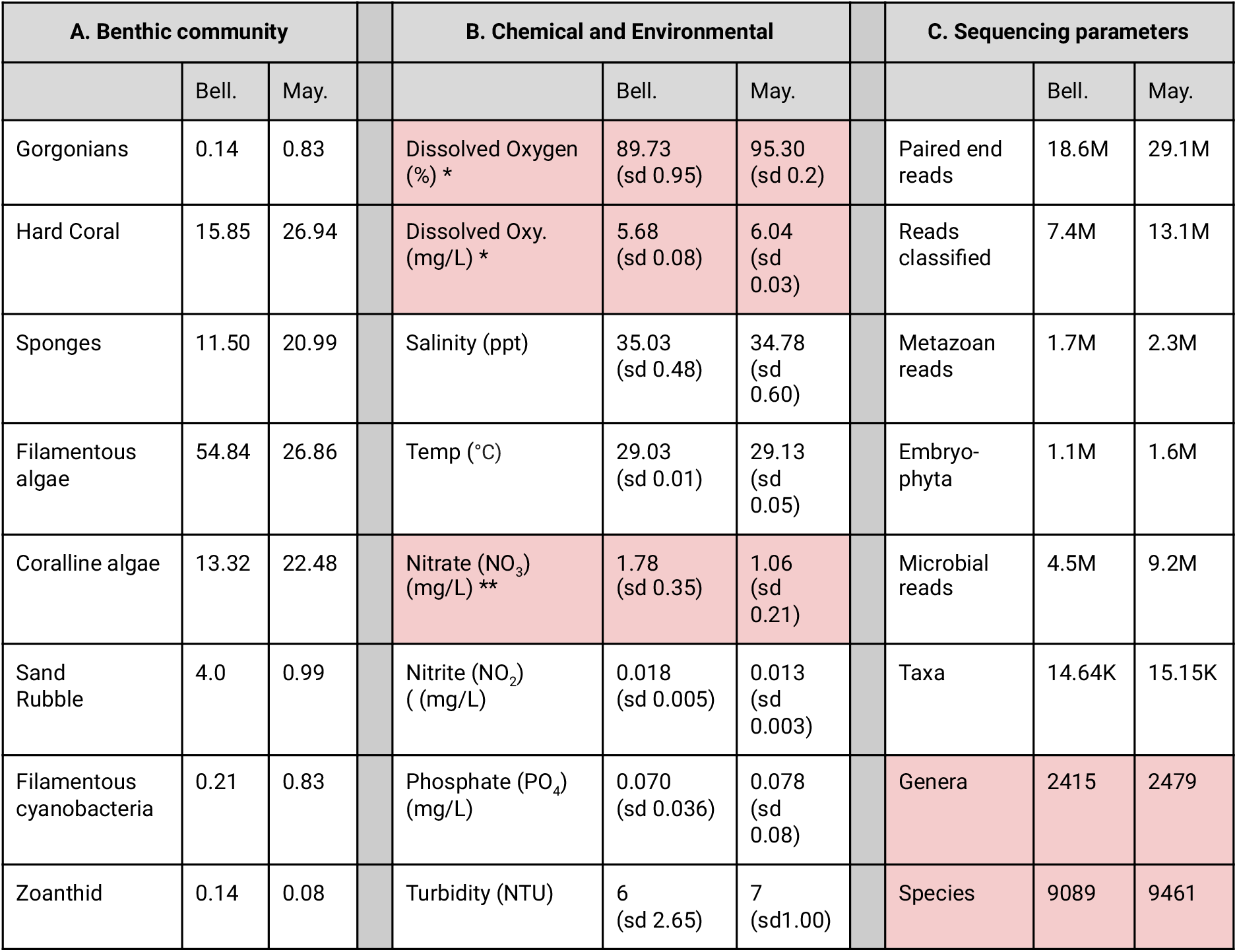
Comparison of the Bellairs and Maycocks sites across different parameters sets. **A**. The benthic composition of the Bellairs and Maycocks Reef based on the percentage cover of major benthic categories on the substratum (from Oxenford and Valles, 2016). **B**. Measurements of several environmental and chemical variables. The * and ** denote signifcance at p < 0.01 with a t-test and Wilcoxon test respectively. **C**. Results of the whole genome shotgun metagenomic sequencing.

| A. Benthic community |  |  | B. Chemical and Environmental |  |  | C. Sequencing parameters |  |  |
| --- | --- | --- | --- | --- | --- | --- | --- | --- |
|  | Bell. | May. |  | Bell. | May. |  | Bell. | May. |
| Gorgonians | 0.14 | 0.83 | Dissolved Oxygen (%) * | 89.73 (sd 0.95) | 95.30 (sd 0.2) | Paired end reads | 18.6M | 29.1M |
| Hard Coral | 15.85 | 26.94 | Dissolved Oxy. (mg/L) * | 5.68 (sd 0.08) | 6.04 (sd 0.03) | Reads classified | 7.4M | 13.1M |
| Sponges | 11.50 | 20.99 | Salinity (ppt) | 35.03 (sd 0.48) | 34.78 (sd 0.60) | Metazoan reads | 1.7M | 2.3M |
| Filamentous algae | 54.84 | 26.86 | Temp (°C) | 29.03 (sd 0.01) | 29.13 (sd 0.05) | Embryo-phyta | 1.1M | 1.6M |
| Coralline algae | 13.32 | 22.48 | Nitrate (NO<sub>3</sub>) (mg/L) ** | 1.78 (sd 0.35) | 1.06 (sd 0.21) | Microbial reads | 4.5M | 9.2M |
| Sand Rubble | 4.0 | 0.99 | Nitrite (NO<sub>2</sub>) (mg/L) | 0.018 (sd 0.005) | 0.013 (sd 0.003) | Taxa | 14.64K | 15.15K |
| Filamentous cyanobacteria | 0.21 | 0.83 | Phosphate (PO<sub>4</sub>) (mg/L) | 0.070 (sd 0.036) | 0.078 (sd 0.08) | Genera | 2415 | 2479 |
| Zoanthid | 0.14 | 0.08 | Turbidity (NTU) | 6 (sd 2.65) | 7 (sd 1.00) | Species | 9089 | 9461 |

**Figure 1.**
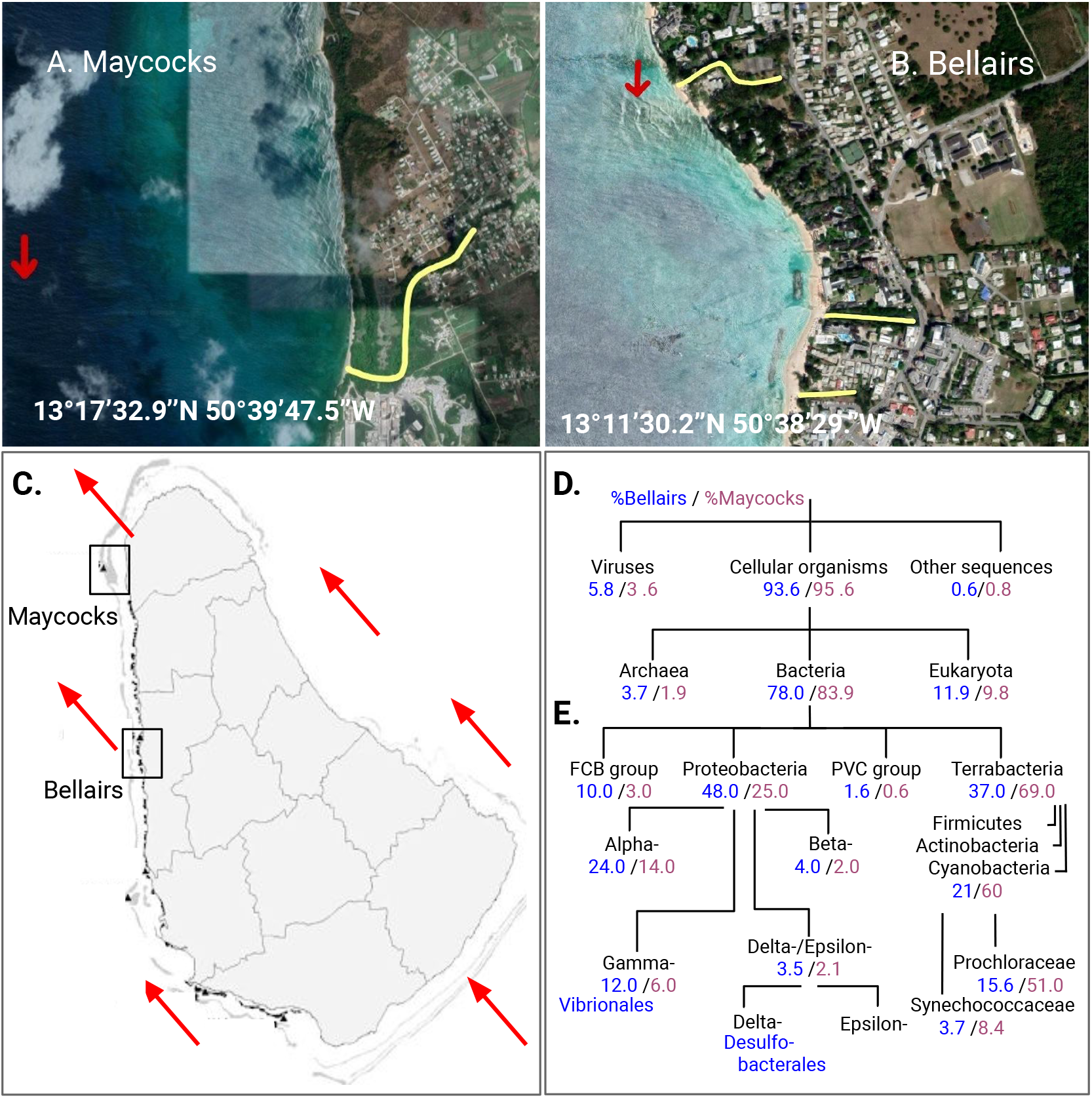
Aerial view of the Maycocks (**A**) and Bellairs (**B**) sites with surrounding areas. Yellow denotes runoffs. Red arrows denote sampling sites. **C**. Map of Barbados with the sea current direction (red). **D**. Percentage of total read counts at the root of the tree of life after removing contaminants. **E**. The subtree of Bacteria.

### Whole genome sequencing of the microbiome of reef water from the two sites

DNA was extracted and whole genome shotgun sequenced (NovaSeq 2 x 150bp, Illumina Inc., **Methods 3**,**4**). The resultant paired-end reads were first processed by our bioinformatic pipeline for read trimming, quality control (**Methods 5**). The Kraken2^38^ and Bracken^39^ packages were then used to align the reads against a compendium of databases from the microbial subset of the NCBI^40^ and Mar marine reference database^41^ in order to estimate taxa abundances (**Methods 6**). Read counts from both sites were mapped to nodes of the tree of life created using the NCBI Taxonomy database^42^ (**Methods 7**).

Reads mapped to non-microbial taxa were removed from the dataset, but we note that analysis of reads mapped to Metazoa, Embryophyta and multicellular fungi identified marine and terrestrial species indigenous to Barbados and ornamental plants and crops grown along the coast (**Supplemental Information 1**). The final microbial metagenomic dataset after filtering and quality control consists of 4.5M and 9.2M reads at Bellairs and Maycocks respectively (**Table 1C**). In total, the profiles identified 2415 (Bellairs, B) and 2479 (Maycocks, M) genera and 9089 (B) and 9461 (M) species. Convergence to these numbers occurs when the read data is downsampled to just one-third of the total dataset, and other measures of ecological richness failed to predict any additional taxa at either site (**Supplemental Information 2**). We built linear models to adjust abundance measures relative to genome size (**Supplemental Information 3**).

Maycocks exhibits a relative enrichment of Bacteria and Chlorophyta whereas Bellairs contains more eukaryotic (primarily fungal) and archaeal taxa (**Figure 1D**). All of these differences are significant (p << 0.01, Pearson’s χ^2^, **Methods 8a**). This global breakdown is consistent with the Tara Oceans 2009-2013 project^8,33,43^ which did not sample in the vicinity of Barbados: ∼81% of all reads were mapped to Bacteria and 3% of all reads as Archaea in both datasets. They diverge for viruses (7.5% Tara vs 4.7% Barbados) and Eukaryota (4.6% Tara vs 10.9% Barbados)^33^.

### Bellairs is enriched for bacteria involved in recycling of organic matter

There was evidence for 6,279 species at Bellairs and 6,459 species at Maycocks across 1,434 distinct bacterial genera, although the distribution in abundances differs significantly between the sites (**Figure 1E, Figure 2A**). Bellairs is enriched in gram negative Proteobacteria (48% B vs 25% M) and the Fibriobacteres-Chlorobi-Bacteroidetes (FCB) group (10% B vs 3% M) where Maycocks has a strong preference for the Terrabacteria group (37% B vs 69% M). These differences are all highly significant (p << 0.01, **Methods 8a-c**). **Figure 2A** (and **Supplemental Figure 8**) establishes that the majority of differentially abundant genera correspond to Proteobacteria and Terrabacteria. Within the PVC group, the Planctomycetes were more prevalent at Bellairs than Maycocks (Kruskal-Wallis KW, p << 0.01, **Methods 8c**) with the genus Rhodopirellula exhibiting differential abundance. Rhodopirellula is a widely distributed marine genus that plays an important role in global carbon and nitrogen cycling^44^. Mariniblastus fucicola and other microbiota associated with coral degradation and algae cover are enriched at Bellairs^45^.

**Figure 2.**
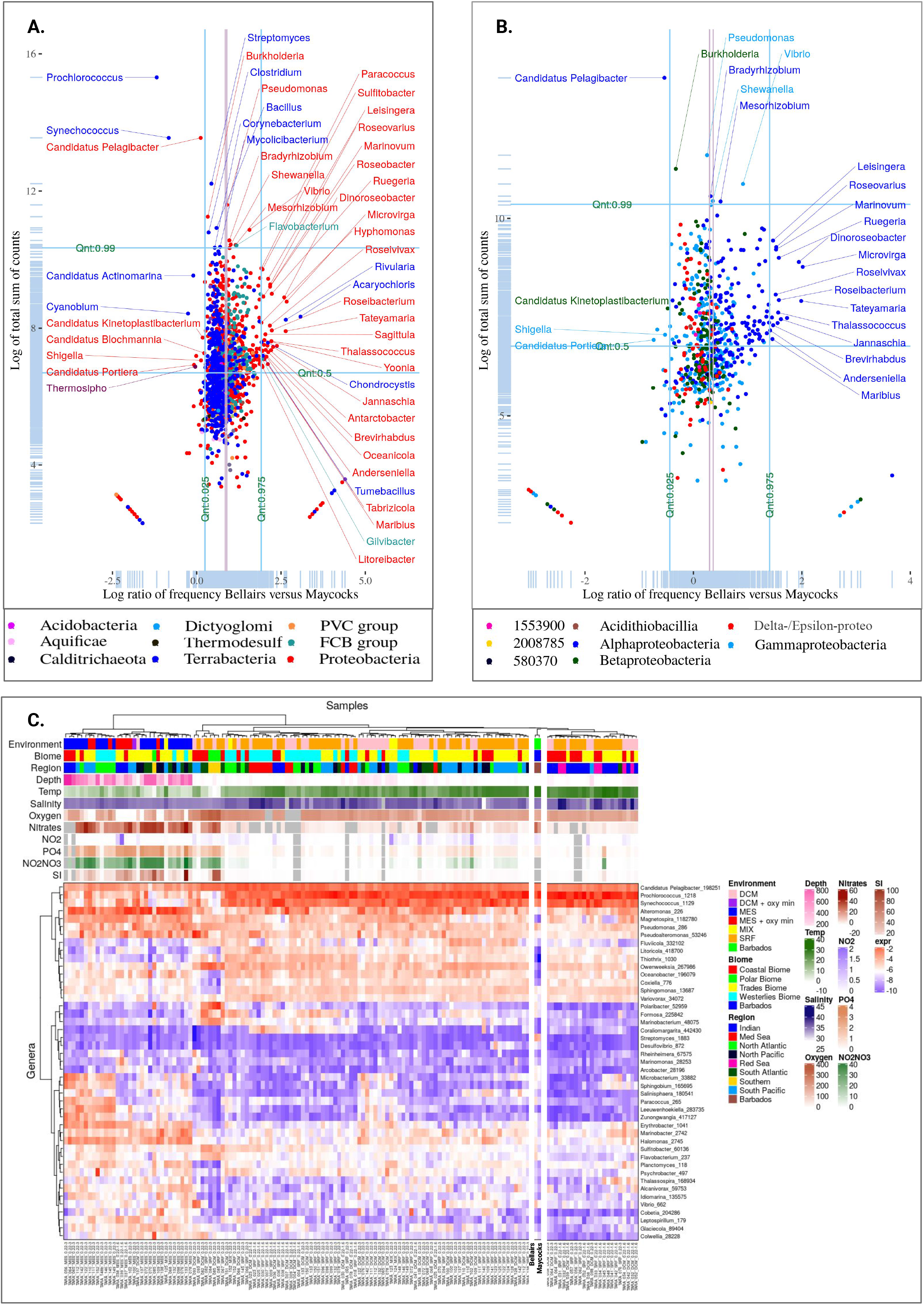
**A.** The log ratio of the fraction of all bacterial reads at Bellairs versus the fraction of all bacterial reads at Maycocks across all genera within the Bacteria superdomain. **B**. Analogous to panel A except here attention is restricted to genera within Proteobacteria. **C**. A heatmap of relative abundance of individual species, and associated measures of ecological diversity, and physio-chemical-hydrographic attributes integrating the Barbadian coral reef profiles with the Sunagawa et al. Tara Oceans data.

With respect to the FCB group, Bacteroidetes are significantly enriched at Bellairs (8.2% B vs 2.8% M, KW, p << 0.01). These gram-negative non-spore forming bacteria are often detected in the gut of animals, and also seawater, where they degrade polymeric organic matter.

Maycocks showed a significant enrichment of the genera Candidatus Actinomarina, ultra small free living species that perform photoheterotrophic metabolism and mirror similar geographic distributions to picocyanobacteria species^46^. However, many other Actinobacteria are commonly found in the soil of marine ecosystems where they play a role in recycling refractory biomaterials and perform nitrogen fixation in symbiosis with plants^47^. Such Actinobacteria are enriched at Bellairs including Illumatobacter coccineus and Arthrobacter sp. LS16.

### Maycocks is highly enriched for phototrophic Terrabacteria

Terrabacteria is the most discordant of all taxa between the two sites (37% B vs 69% M). The vast majority of the Maycocks Terrabacteria reads are mapped to Cyanobacteria (60 of 69%; **Figure 1E**). Cyanobacteria are phototrophic bacteria that synthesize organic compounds from carbon dioxide, and produce oxygen as a by-product. They are extremely abundant in nutrient-poor warm waters. In our data, the autotrophic genera Prochlorococcus and Synechococcus account for over 98% of all Cyanobacteria identified in our study (**Figure 2A**), consistent with the oligotrophic nature of the Barbadian marine environment^48^. Prochlorococcus is typically more prevalent in nutrient poor water while Synechococcus is more prevalent in eutrophic water and coastal plumes of rivers, likely due to increased nitrate and phosphate levels^49^. This is consistent with our data where we observe a 3.7:1 Prochlorococcus to Synechococcus ratio at Bellairs, but a 5.3:1 ratio at Maycocks. A study of the coral reef ecosystems in the Caribbean waters of Curacao also reported a high percentage of Cyanobacteria (30-43%), well beyond levels observed in the north Atlantic Ocean, Mediterranean Sea and Pacific Ocean^50^. Moreover, a recent study of northern Caribbean reef systems also highlighted high abundance of Cyanobacteria among their sites with a relative abundance ranging from 13.2% to 29.7% of all bacterial and archaeal phyla, again primarily due to the presence of Prochlorococcus and Synechococcus^25^. Our observed frequencies for Cyanobacteria at Maycocks are significantly more extreme.

The remainder of Terrabacteria show enrichment at Bellairs including the Firmicutes, an extremely diverse group of endospore-forming bacteria able to survive in extreme environments and often found in nutrient-rich environments including marine sediment^51^ (**Supplemental Figure 9**). Several genera were more abundant at Bellairs including the Tumebacillus (gram-positive spore forming sulphur-oxidising bacteria that have been isolated from diverse environments such as algal scum, freshwater, soil and mangroves^52^ and within the skeletal mucus of Caribbean coral porites^53^) and the Fictibacillus (aerobic bacteria of which some species possess protease-producing abilities that play a major role in the biodegradation of corals^54^).

### Bellairs is enriched for Alphaproteobacteria associated with algae and ocean sediment

Alpha- and Gamma-proteobacteria are believed to contribute the majority of bacterioplankton to ocean waters, an observation consistent with the relative abundances in our data (Alphaproteobacteria 24% Bellairs vs 14% Maycocks, Gammaproteobacteria 12% Bellairs versus 6% Maycocks; **Figures 1E**). The shift towards Bellairs is significant in both cases (KW and Dunn’s test, p <<0.01, **Methods 8c, Figure 2B**). Rhodobacterales can utilize various organic and inorganic compounds and carry out sulfur oxidation, aerobic anoxygenic photosynthesis, carbon monoxide oxidation and the production of secondary metabolites; there is evidence that members of Rhodobacterales correlate with the sedimentary setting^55^ (**Supplemental Figure 10**). Although some Rhodobacteraceae bacterium strains were highly abundant and biased towards Maycocks, the vast majority of genera showed a general trend towards the Bellairs site (KW test, p << 0.01) highlighting genera Dinoroseobacter, Tateyamaria and Jannaschia. Dinorosebacter are aerobic anoxygenic phototrophic bacteria, known to be highly abundant in marine turf algae. Some Dinoroseobacter species form epibiotic relationships with red tide dinoflagellates^56^. Tateyamaria is a genus of marine gram-negative aerobic bacteria isolated from coastal marine environments. Tateyamaria have been identified as components of soft corals and coralline alga microbiomes. Within the alga microbiome, Tateyamaria species are able to survive and increase abundances under acidification conditions^57^. Jannaschia are aerobic anoxygenic phototrophic bacteria. Some species play a role in transport and nitrate reduction^58^. Other species appear to be part of microbial communities associated with corals^59^.

Conversely, the oligotrophic genus Candidatus Pelagibacter is significantly shifted towards Maycocks (KW test, p << 0.01). These small free-living heterotrophic species, which are one of the most abundant genera identified in our study, are usually found thriving in low-nutrient environments, which play a significant role in carbon cycling, feeding on dissolved organic carbon and nitrogen^60,61^. The species Candidatus P. ubique belongs to the SAR11 clade ubiquitous in the world’s oceans ^61^.

### Bellairs is enriched for pathogens of many dimensions of the reef biosystem

Within Gammaproteobacteria, the genus Vibrio was one of the most highly abundant genera in our dataset and exhibits a strong skew towards Bellairs (KW, p << 0.01) involving many species including corallilyticus, tubiashii, harveyi, astriarenae, nigripulchritudo, and ponticus (**Supplemental Figure 11**). Vibrio are gram-negative motile bacteria commonly found in marine environments and are facultative anaerobes, capable of producing ATP by aerobic respiration if oxygen is present, but also able to switch to fermentation. Some Vibrio species play a significant causative role in coral diseases and disrupt corals symbiotic relationship with zooxanthellae. There are indications that high nutrient levels promote pathogenic bacteria including Vibrio spp. to dominate in healthy coral reef systems^62^. In our data, there was evidence for several Vibrio species at Bellairs including V. corallilyticus (implicated in white band syndrome) and V. harveyi (linked to yellow spot syndrome but also present in healthy corals albeit less frequently^63^), V. tubiashii (implicated in shellfish vibriosis and may also be a virulence factor in diseases of scleractinian corals, where it plays a role in photoinactivation of the coral^64^), V. astriarenae (a generalist in many reef systems^65^), V. nigripulchritudo (a shrimp pathogen with major impact on farms in Japan and New Caledonia^66^), and V. ponticus (a fish pathogen^67^). Photobacterium, also a genus of Vibrionales, is common in marine environments and can survive in both aerobic and anaerobic environments. P. damselae, which was more abundant at Bellairs, is a well-studied pathogen of marine organisms including fish and has made significant negative financial impact on fisheries world-wide ^68^. Two species of the genus Acinetobacter within Pseudomonadales are enriched at the Bellairs site. Acinetobacter is a gram-negative genus which plays an important role in the mineralization of aromatic compounds within soil including marine systems, and many species can reduce nitrates to nitrites^69^. Our study highlights A. schindeleri and A. indicus, both emerging opportunistic human pathogens that can survive in many environments^70^.

In contrast, the most differentially abundant at the Maycocks site are several unclassified Gammaproteobacteria including members of the SAR86 clade, which are globally abundant planktonic bacteria^71^.

### Bellairs is enriched for sulfate reducing Deltaproteobacteria

The Delta-Epsilon subdivision displays a dramatic split in their preference between the two sites (**Supplemental Figure 12**). Epsilon members are skewed towards Maycocks (KW test, p<<0.01) but lie close to the 95% confidence interval for the global mean (red bars). The vast majority of Deltaproteobacteria are skewed towards Bellairs including the Desulfomicrobium, a genus of sulfate reducing bacteria that thrive in marine anoxic environments and interfaces such as microbial mats^72^ including the species D. orale which has been previously identified in Atlantic coastal marine waters^73^. Several SAR324 species are highly abundant with a preference for Maycocks. They have a versatile metabolism with heterotrophic carbon utilization capabilities and are implicated in sulfur oxidation^74^.

### Maycocks is a lithotrophic hotspot and Bellairs is similar to other warm ocean waters

We compared our profiles with the Tara Oceans effort which generated 243 samples from 68 locations in epipelagic and mesopelagic waters restricted to only bacterial taxa (n=139, **Methods 9**)^33^. **Figure 2C** directly compares our results with Sunagawa et al.^33^ across several types of variables including measures of ecological diversity, physio-chemical-hydrographic attributes and relative abundances of individual species. Here we used unsupervised clustering to investigate how our sites compare across the global sites sampled by Tara Oceans. Bellairs and Maycocks co-cluster with other samples harvested from the surface and deep chlorophyll maximum layers (SRF, DCM respectively, **Figure 2C**) of the trade and coastal biomes. This is consistent with the finding from Sunagawa et al. that depth is the single most important factor that determines species abundance as it explains 75% of variation via Principal Coordinate Analysis. Both Bellairs and Maycocks are closest to Red Sea and Indian Ocean samples. Although concentrations of nitrates NO_2_ and PO_4_ at Maycocks are similar to levels observed in samples it co-clusters with, nitrate levels of Bellairs are higher and more similar to mesopelagic layers (MES in **Figure 2C**).

Abundance levels across the selected genera are generally consistent with the Tara Ocean data. The right subtree of **Figure 2C** is enriched for autotrophs consistent with the fact that Barbados is a hot climate with oligotrophic waters. However, several genera were under-represented at both locations in comparison to Tara Oceans. This includes Magnetospira, Litoricola, Thiothrix, Owenweeksia, Oceanobacter and Coxeilla. Conversely, Streptomyces at Bellairs was an outlier exhibiting an abundance higher than all other samples.

### Both sites exhibit high levels of methane producing anaerobic Archaea

In total, there was evidence of 440 species at Bellairs and 442 species at Maycocks across 122 distinct archaeal genera. A high percentage of all archaeal reads were mapped to Euryarchaeota at both sites, although there is a significantly higher fraction at Maycocks (63% B vs 82% M; **Figure 3A**). For both sites, the second highest fraction of archaeal reads were mapped to the TACK (Thaumarchaeota, the ancestor of the Cren-, Cand. Bathy- and Cand. Kor-archaeota) group, although here there is a significantly higher fraction at Bellairs (32% B versus 13% M). These differences are all significant (p << 0.01, **Methods 8**).

**Figure 3.**
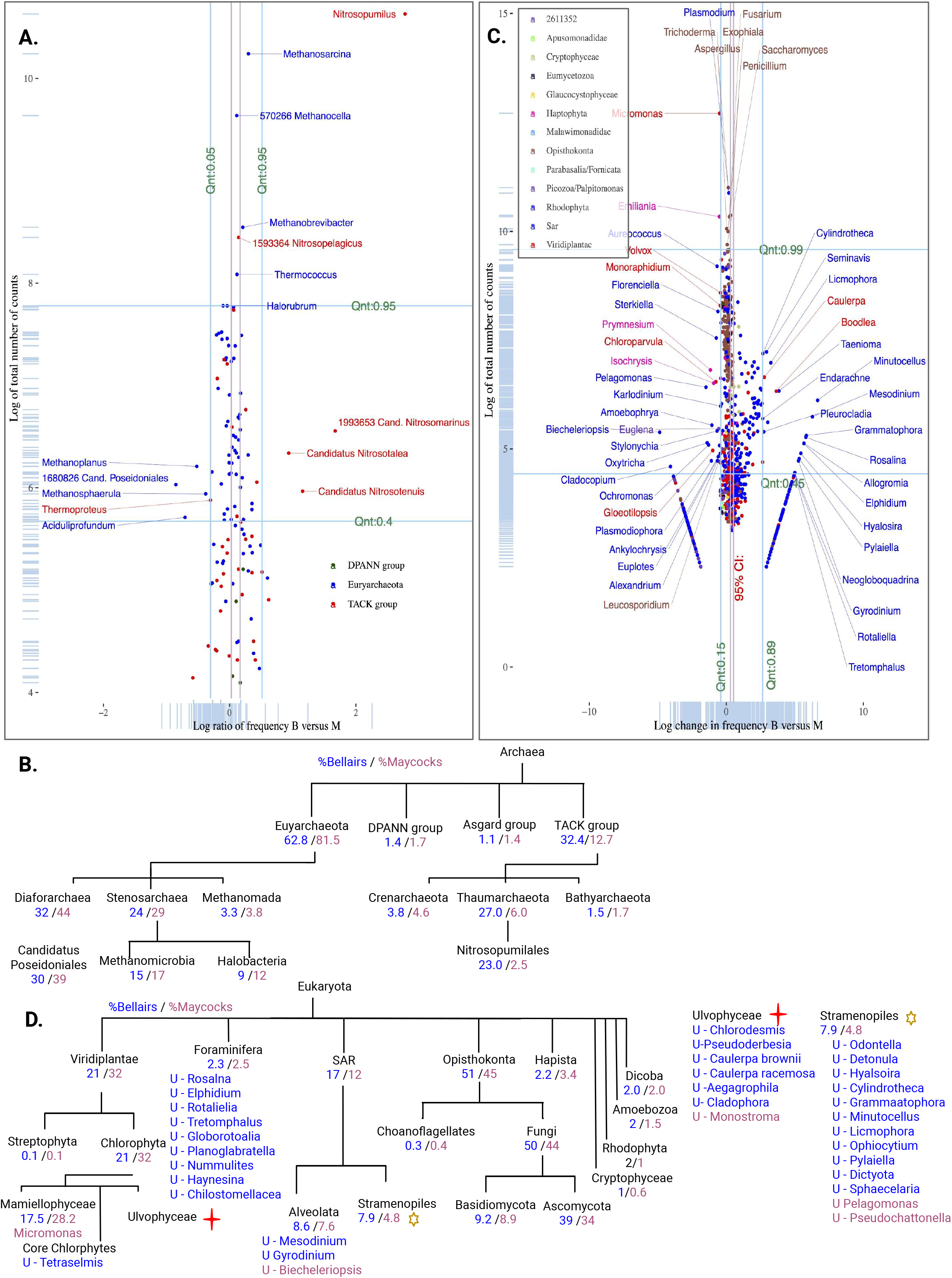
**A.** Analogous to the plots 2A,B but here focus is restricted to Eukaryota. **B**. Percentage of total read counts for abundant sub-taxa of Eukaryota. **C**. The scatterplot for Archaea. **D**. Percentage of total read counts for abundant sub-taxa of Archaea.

Several genera from Methanomada groups are highly abundant in equal proportions at both sites (**Figure 3C**). The Methanosarcina genus contains anaerobic methanogens that conduct methanogenesis in diverse environments throughout the world including seawater and in coral rubble^75^. Methanobrevibacter is a genus of methane producing anaerobic archaea. Some known species of this genus inhabit animal intestinal tracts, decaying plants and sewage and are used as indicators of faecal pollution in coastal waters^76^. Methanocella species are known soil and sediment archaeons^77^. Several genera however including Methanoplanus and Methanoshpaerula have significantly higher relative abundance at Maycocks. This genus contains methanogenic species that are known to play endosymbiotic roles with marine ciliates^78^ and other species associated with sponges in anoxic environments^79^.

### Photoheterotrophic euryarchaeota are highly enriched at Maycocks

The majority of Euryarchaeota reads at Maycocks are mapped to Diaforarchae with smaller amounts mapping to the Stenosarchaea group (24% B vs 29% M), and Methanomoda (**Figure 3 A**,**C**). Species from the Candidatus Poseidoniales order are some of the most abundant planktonic archaeons in ocean surface waters^80^, consistent with the elevated relative abundances in our data. The ancestor of Cand. Poseidoniales is a motile photoheterotroph, capable of degrading proteins and lipids, although there is genus-specific lifestyle and niche partitioning^80^.

### Bellairs is enriched for thaumarchaeota involved in nitrification

The majority of Thaumarchaeota reads at Bellairs are mapped to the Nitrosopumilales order (23% B versus 3% M, Dunn’s test, p << 0.01; **Figure 3C**). Nitrosopumilus are ammonia-oxidizing archaeons commonly found in marine environments^81^. The chemolithoautotrophic Nitrosopumilus maritimus species, which is enriched at the Bellairs site, has been established as a dominant contributor to nitrification^75^. Several additional Nitrosopumilales genera are also differentially identified between Bellairs and Maycocks including Candidatus Nitrosopelagicus, a planktonic pelagic ammonia-oxidizing thaumarchaeon involved in nitrogen and carbon fixation in marine environments^82^.

### Maycocks is highly enriched in photosynthetic autotrophic Chlorophyta

The two sites identified a similar percentage of Eukaryota of which approximately half are Opisthokonta, specifically Fungi (**Figure 3B**). Although several “true yeasts” saccaryomyceta are highly abundant, we did not observe a preference for either site. Viridiplantae exhibits an enrichment at Maycocks (21% B vs 32% M) with 99% of these reads mapping to the autotrophic green algae Chlorophyta (**Figure 3D**). This includes Micromonas pusilla and commoda which are photosynthetic picoeukaryotes known to thrive globally in tropical marine environments^83^ and play a key role in the primary production within the euphotic zone^84^. These findings are consistent with a recent Caribbean study highlighting Micromonas^85^.

Boodlea composita, which is abundant at the Bellairs site, is a common green algae in coastal marine environments^86^. These macroalgae are capable of blooms forming dense turf or mats in nutrient rich waters. B. composita growths have been observed to smother coral colonies^87^. Species of the genus Caulerpa were present only at Bellairs. Overall the three species had low abundance, although C. brownii was witnessed by just over 200 reads. Caulerpa is a genus of nitrophilic macroalgae that commonly inhabit tropical marine environments including Caribbean coral reefs^88^. Like B. composita, some species can bloom creating carpets which smother coral colonies^89^. Caulerpa species assimilate nutrients from sediment and grow successfully in habitats with anthropogenic disturbance such as waste and stormwater^90^.

### Maycocks is enriched for organisms involved in primary production and regulation of the micro-planktonic community

The Stramenopiles-Alveolates-Rhizaria (SAR) supergroup, which we winnowed for single cellular organisms, exhibits broad heterogeneity, contributing outlying species and genera at both sites (**Figure 3D, Supplemental Figure 13**). Several species of the SAR supergroup species are more highly abundant at Maycocks. Genus Pelagomonas is monotypic containing only the picoplankton P. calceolata, a tiny photosynthetic flagellated alga that contributes to primary production in marine environments^91^. Biecheleriopsis contains small marine phototrophic planktonic dinoflagellates that play a role as primary producers and symbiotic partners^92^. Karlodinium are phytoplanktonic coastal dinoflagellate mixotrophics that rely on both photosynthesis and phagotrophy and can cause toxic algal blooms when nutrients are scarce within their environments^93^. Euglena species are heterotrophic primary producers within marine ecosystems^94^. Alexandrium is a genus of planktonic dinoflagellates that contribute to primary production^95^.

Other species and genera biased towards Maycocks are regulators of the micro-planktonic community or have symbiotic roles with coral and sponges. These include for example the Stylonychia^96^, Amoebophrya (marine parasitic dinoflagellates that inhabit coastal water^97^), the golden algae Ochromonas (small unicellular mixotrophic flagellates that play a key role in regulating bacterial abundance^98^), the dinoflagellates of Cladocopium (that establish endosymbiosis with cnidarian species such as coral^99^), Cladocopium species (dominant symbionts of some stony corals within the Caribbean including Barbados^100^), and Euplotes (filter-feeding ciliates linked to the ingestion of coral tissue^101^ (**Figure 3C**).

### Bellairs is enriched for benthic epiphytes growing on coral and algae

Many species of Stramenopiles are observed uniquely at Bellairs (**Figure 3D** denoted by orange star, 11 of all 34 uniquely identified organisms identified at Bellairs, hypergeometric binned by eukaryotic phyta, p << 0.01). This is not likely due to the depth of sequencing alone given that Maycocks received 1.57 fold more reads than Bellairs and given our downsampling experiments described earlier.

Several marine diatoms were identified including Licmophora (common epiphytes abundant on substrates such as filamentous algae or corals where they can form mats which smother coral^102^); Cylindrotheca (ubiquitous in coastal areas worldwide that utilize latent nitrogen and phosphorus nutrients in the environment^103^); Seminavis (often associated with seaweed living within coral reef ecosystems^104^); Minutocellus (specifically species M. polymorphus, a symbiont of benthic Foraminifera^105^ found to be codominant with brown tide, which thrives in eutrophic marine environments^106^); Hyalosira (known to attach to seaweeds in intertidal environments^107^); and, Grammatophora (an epiphytic marine genus of diatoms found in coastal marine environments^108^). Furthermore, Bellairs identifies members of the genus Gyrodinium (marine heterotrophic dinoflagellates which prey on diatoms and can cause red tides^109^), and Pleurocladia (benthic brown alga epiphytes of macroalgae).

### Bellairs exclusively has low levels of Foraminifera

Analogous to the Stramenopiles, the Foraminifera contribute a surprising number of uniquely identified genera (9 of 34; p << 0.01, hypergeometric binned by clades directly from the root of Eukaryota; **Figure 3D**). There was no evidence of a Foraminifera taxon at Maycocks. Foraminifera are single-celled shelled protists and recognized as one of the most abundant groups of microorganisms in the shallow marine waters. The fact that their size range (100µm-20cm) is well beyond our filtered range may explain the paucity of reads for these taxa. Some amoeboid protists are often benthic or live in the sea sediment; at least 40 morphospecies are planktonic and form symbiotic relationships with marine algae. They are sensitive to the subtle changes in the ambient environment and species are known to survive and increase in numbers in polluted areas^110^. Planktonic foraminifera play an important role in the carbonate pump, contributing up to 50% of the total carbonate in the ocean sediment^111^. Genera Neogloboquadrina, Globorotalia, Planoglbratella, Elphidium, Rosalina, Allogromia and Rotaliella are all recognized as planktic. Rosalina, for example, is known to attach to seaweed and other marine benthic surfaces within shallow environments and can also be found unattached within sediment^112^. Species such as R. leei are able to thrive in ecologically stressed environments^113^.

### Other protists and single cell algae support primary production at Maycocks

With respect to the remaining subclades of Eukaryota, the Haptophyta are biased towards Maycocks (Dunn’s test, p < 0.01) and contribute several outlying genera including the coccolithophorid genus Emilliania. The main species within this genus is E. huxleyi, a unicellular photosynthetic eukaryote that is a key contributor to the oceanic carbon cycle via calcification, photosynthesis and export of inorganic matter to the oceans’ interior^114^. The planktonic unicellular flagellates Prymnesium are also more abundant at Maycocks.

### Cyanophages are highly enriched at Maycocks, a site highly enriched for cyanobacteria

Metagenomic analysis of marine viruses was investigated within the context of the Tara Oceans Project^115^ but these efforts used sequencing and bioinformatics platforms specific for viromes. Our investigation here is limited, since we selected for organisms between 0.22 µm and 3 µm. Nevertheless, 5.8% (B) and 3.6% (M) of all reads mapped to viruses (**Supplemental Information 4**). Consistent with the strong preference for Cyanobacteria (incl. Prochlorococcus and Synechococcus) at Maycocks (26% B vs 62% M), there is a comparably strong preference for cyanophages at Maycocks (KW and Dunn’s test for Caudovirales, both p << 0.01). Other well-established phages of Prochlorococcus and/or Synechococcus^116^ including Myo-, Sipho- and Podo-viruses are also identified in our data. In contrast, the small heterotrophic Pelagibacter, a member of the ubiquitous SAR11 clade, is highly enriched at Maycocks, however Podovirus phages of Pelagibacter are systematically shifted towards Bellairs (KW, Dunn’s test, p<<0.01), suggesting enrichment of the Podoviruses at Bellairs.

## Discussion

The two sites of the Barbadian coral reef system studied here differ in environmental and ecological attributes. This includes differences in their endogenous benthic communities including hard corals, coralline and sponges. Although both have suffered under the same global stressors, there is likely a differential in the effects of local stressors. For example, eutrophication has long been postulated as a major negative influence on the health of Barbadian reefs which lie close to urban areas and coastal run-offs^16^ and corresponding studies of the macrobenthic community have noted decay in almost every dimension of such ecosystems^17^. This is the first study to use whole genome shotgun sequencing of the Barbadian reef water microbiome. The sequence data identifies over 9,000 species across the tree of life at both sites, and the composition of both communities appears in concordance with previous findings regarding coral reef health in Barbados. Although our study here is limited in scope, our samples do share many similarities with other ocean sites including the Tara Ocean project and several Caribbean specific projects, suggesting that the sequencing has captured many relevant features of the ecosystems.

Although it is infeasible to quantitatively estimate the physiological relevance of the observed differences in relative frequencies of taxa, there are striking and consistent differences between Bellairs and Maycocks. Maycocks is strongly concentrated with phototrophs and is a clear outlier in this dimension when compared against data from the Tara Ocean project. The levels of Prochlorococcus and Synechococcus are only matched by a study of nearby Curacao, suggesting that the southern Caribbean constitutes a cyanobacterial hotspot. The relative abundance of photosynthetic organisms other than the cyanobacteria are also significantly higher at Maycocks than Bellairs including genera Candidatus Pelagibacter, Candidatus Poseidoniales, Micromonas and the Emilliania.

In comparison, Bellairs is enriched for copiotrophs, macroalgal symbionts and marine-related disease-bearing organisms from taxa scattered across the tree of life. For Bacteria alone, this includes species from the Firmicutes (found in nutrient-rich extreme environments), species from the Planctomycetes (with established roles in coral degradation where turf algae is dominant), many Rhodobacterales species (that utilize compounds in a nutrient rich environment for the production of secondary metabolites), at least six genera of Dinoroseobacter (commonly found in marine turf algae and which participate in nitrate reduction), the Desulfomicrobium (sulfate-reducing bacteria that thrive in marine anoxic environments and in interfaces such as microbial mats), the Acinetobacter (associated with mineralization of aromatic compounds), and several species of the genus Vibrio (well implicated in marine diseases including white band and yellow spot syndrome). The Archaea highlight the Nitrosopumilales genera (pelagic ammonia-oxidizing thaumarchaeon involved in nitrification and carbon fixation). The Eukaryota include the Boodlea (nitrophilic macroalgae species capable of rapid growth in nutrient rich waters), in addition to many diatoms and dinoflagellates (Licmophora, Cylindrotheca, Seminavis, Gyrodinium, Pleurocladia, Minutocellus). These all have established roles in marine coral reef settings including the ability to form algal mats which smother coral.

Our data indicates a shift from coral to macroalgae between Bellairs and Maycocks, an observation with potential importance for the design and feasibility of coral rehabilitation projects around the island. More generally, broader longitudinal studies would provide more information regarding the rate and direction of change, and may perhaps identify ecological markers suitable for diagnostic, prognostic or predictive purposes when integrated with macrobenthic monitoring and international oceanic profiling projects.

## Methods

Analyses here were carried out in R version 4.03^117^.

### 1. Sample collection and preparation

Samples were collected during the afternoon from two locations along the west coast of Barbados; Folkestone Marine Reserve (13°11’30.2”N 59°38’29.2”W) and Hangman’s Bay (13°17’32.9”N 59°39’47.5”W) on January 30th and 31st 2018 respectively. At the time of sampling the ocean current was flowing in a northwestern direction between the two sample days^118^. Sea water was collected from the Bellairs reef site located in Folkestone Marine Reserve just below the surface. Seawater was collected at 1 metre above the Maycock reef located in Hangman’s Bay using SCUBA and a boat for reef access. All samples were collected using 7L acid-washed bottles between 1-3pm in the afternoon. The samples were transported back to shore and immediately passed through a 3µm pore polycarbonate membrane filter, followed by 0.22 µm pore Sterivex filter, in both cases using a peristaltic pump (7” of mercury). Organisms captured by the 3µm filter were discarded. RNA later was added to the 0.22µm filter and stored at −80°C. We therefore expected that organisms smaller than 0.22µm are removed from the analysis. The range of organism size is consistent with the sample collection specifications of the Tara Oceans project^33,34^. Surface sea water temperature at time of collection was estimated using archived data and images from the NOAA Coral Reef Watch^119^.

### 2. Nutrient analysis and estimates of microbial content in the seawater samples

Water measurements were taken using an EXO2 Multiparameter Sonde, 20m apart at a depth of 5m at both sampling sites. This analysis was conducted on September 4th 2020. Seawater temperature, salinity and dissolved oxygen information was collected (3 sample readings per location). Sea water samples were also collected to conduct analysis of nitrates, nitrites, phosphates and turbidity. These analyzes were conducted by the Centre for Resource Management and Environmental Studies (CERMES, Barbados). Nitrate and nitrite concentration was determined using the cadmium reduction method, phosphate concentration by the ascorbic acid method, while turbidity was determined using a turbidity meter.

Direct counts of Bacteria, Archaea and Virus-like particles were generated using methods described previously^120^. Aliquots of the seawater samples used for the metagenomics were stored at −80°C in acid-washed two 60ml polycarbonate bottles with TE-glycerol to each bottle, they were then immediately stored at −80°C. At time of microscopy, samples were unfrozen and aliquots of 6 ml per sample were fixed with Electron Microscopy Sciences 4% Paraformaldehyde (5025999, Fisher Scientific), stained with SYBR™ Gold Nucleic Acid Gel Stain (S11494, ThermoFisher) and centrifuged for 3 hours. The samples were placed onto 0.02 µm Anodisc inorganic filters (WHA68096002, Millipore Sigma) and mounted on glass slides. Counting was done using epifluorescence microscopy. Images were captured on a Nikon Ti inverted microscope equipped with metal halide lamp (Nikon), GFP filter (480/40ex, 520/75em), and Photometrics Prime BSI camera using NIS Elements software. Images were then preprocessed using FIJI ImageJ.

### 3. Nucleic acid extraction

DNA was extracted from the Sterivex filter using the DNEASY PowerWater Kit (14900, Qiagen Inc.) with an additional 37°C Incubation step. The Sterivex filter was thawed, unfolded and carefully placed into a PowerWater bead tube. To initiate cell lysis, 1 ml of a buffer composed of guanidine thiocyanate (PW1) was preheated to 55°C for 10 minutes and added to the bead tube. The bead tube was placed horizontally to incubate at 65°C for 10 minutes. The bead tube was then vortexed at 3000 RPM for 5 minutes. After vortexing, the bead tube was centrifuged for 1 minute at 3,000g. Once centrifuged, 600-650 µl of supernatant was transferred to a new 2 ml collection tube. Then, 1µl of RNAse A was added and the tube was incubated at 37 °C for 30 minutes. After incubation the tube was centrifuged at 13,000g for 1 minute; the supernatant was then transferred to a new 2 ml collection tube and 200 µl of an IRS solution (PW2) was added. The tube was then vortexed briefly followed by incubation at 4°C for 5 minutes, then centrifuged for 1 minute at 13000g. The supernatant was then transferred to a new 2ml collection tube; 650 µl of a high concentrated salt solution (PW3) was preheated to 55°C and added; the tube was then vortexed briefly. 650 µl of the supernatant was loaded onto a MB Spin Column Filter and centrifuged at 13000g for 1 minute. The flow-through was discarded and centrifugation was repeated until all of the supernatant was processed. The MB Spin Column was placed into a new 2 ml collection tube; 650 µl of an alcohol-based solution (PW4) was added. The tube was then centrifuged for 1 minute at 13000g. The flow-through was discarded and 650 µl of ethanol (PW5) was added. The tube was then centrifuged at 13000g for 1 minute; the flow-through was discarded and the tube was again centrifuged for an additional 2 minutes. The MB Spin Column was then placed into a new 1.5 ml tube lid removed); 50 µl of an elution buffer (PW6) was added and the tube was left to sit for 2 minutes. The tube was centrifuged at 13000g for 1 minute. The supernatant was then transferred to a new 1.5 ml tube (with lid) and stored at −80°C.

### 4. DNA sequencing

In preparation for sequencing the two DNA extracted samples were thawed and resuspended in 10 mM Tris-HCl pH 8.0 with 0.1mM of EDTA. Next generation DNA-level sequencing was performed on the two samples at the McGill University and Genome Quebec Centre on the NovaSeq PE 150 platform generating 2 x 150bp paired-end reads. Library preparation was performed by Genome Quebec based on the following protocol. gDNA was quantified using Quant-iT™ PicoGreen dsDNA Assay Kit (P11496, Life Technologies Inc). Libraries were generated using NEBNext Ultra II DNA Library Prep Kit for Illumina (E7103, New England Biolabs). The IDT Unique Dual Index adapters and universal primers used were [AGATCGGAAGAGCACACGTCTGAACTCCAGTCAC][AGATCGGAAGAGCGTCGTGTAGGGAAAGA GTGT]. Size selection of libraries at 360bp was performed using SparQ beads (Qiagen Inc.). Libraries were quantified using Kapa Illumina GA with Revised Primers-SYBR Fast Universal kit (Kapa Biosystems Inc.). Average size fragment was determined using a LabChip GX (PerkinElmer Inc.) instrument.

### 5. Quality control of sequencing results

The total number of raw paired end reads (2 x 150bp) returned by NovaSeq was 19 and 30.5 million at Bellairs and Maycocks respectively. The quality of reads was first assessed using FastQC version 0.11.5^121^, which provides several metrics for assessing the overall quality of reads including GC bias, sequence quality, base sequence content, base N content, sequence length distribution, sequence duplication levels, overrepresented sequences, the adapter, the distribution of observed k-mers, and the reliability and quality of all bases at each position along a read. Illumina adapters from the paired-end reads were trimmed using Trimmomatic version 0.38 with parameters ‘ILLUMINACLIP:NovaSeq.fa:2:30:10 LEADING:3 TRAILING:3 SLIDINGWINDOW:4:15 and MINLEN:36’^122^. Trimmomatic also filtered out 2.1M reads which were either found to be unpaired or had a length less than 36bp. Quality of the reads was assessed post-trimming using FastQC. Unpaired, short or poor quality reads were removed (4.1% and 4.5% of all reads for Bellairs and Maycocks, respectively).

### 6. Read alignment & taxa classification via Kraken2/Bracken

We used Kraken2^38^ to align and classify the observed sequencing reads against a range of genomes. Kraken2 compares the distribution of k-mers in a query sequence, which is a read (or pair-end read after pre-processing), against the k-mer distribution of genomes in a database of target taxa. For each query sequence, Kraken2 identifies the least common ancestor (LCA) of these taxa in our target genomes which include the plasmid, viral, protozoa, plant, UniVec, env_nr, nr, bacteria, archaea and fungi NCBI downloads and the MAR reference database for marine metagenomics^41^. Bracken^39^ was then applied to the taxonomic assignments made by Kraken2. Bracken uses the Kraken2 assignments in addition to information about the genomes themselves to better estimate abundance at the species level, the genus level, or above. Bracken was used here with a k-mer length of 35 and a read length of 100. Bracken discards reads based on a given threshold. Taxa with total reads less than the set threshold of 10 reads were discarded.

### 7. Taxonomic analysis

Our in-house analysis made use of the NCBI Taxonomy database (version March 31, 2020)^40^. After importing the Taxonomy database into R, we mapped read counts for both sites on the nodes of the tree. Reads mapped to non-single cell taxa were removed from the analysis (**Supplemental Information 1**). The data was examined for ecological richness/diversity and adjusted to handle zero counts (**Supplemental Information 2**) and for genome size (**Supplemental Information 3**).

### 8. Compositional Data and Comparative analyses

Our metagenomic data is compositional in nature^123-125^ and we have attempted to follow so-called CoDa (Compositional Data) best-practice guidelines^124,125^.

8a. A standard Pearson’s χ^2^ test was used to test for statistically significant differences between two multinomial distributions with p-value thresholds depending on the context (chisq.test R function). Here the null distribution is that both sites were generated by a single multinomial distribution.

8b. A two-sided binomial test was used to test for statistically significant differences between the observed marginal frequency(for a component of the multinomial) versus its frequency according to the multinomial distribution (function binom.test R function).

8c. A Kruskal-Wallis test was used to test for statistically significant differences between two or more groups (taxa, described as multinomial distributions derived from our observed data; kruskal.test R function). The null hypothesis is that both sites were generated by the same distribution. When the alternative hypothesis is accepted, it implies that at least one group stochastically dominates another group. Significance was estimated using a p-value threshold. Dunn’s test was used to identify which taxa is stochastically dominant.

### 9. Comparison with the Tara Oceans marine metagenomic samples

The metagenomic sequences and the associated metadata were downloaded from the companion website of Sunagawa and colleagues^33^. Samples that were not filtered for organisms in the range of 0.22µm and 3µm were excluded from our analyses. With respect to the metadata, we primarily made use of Tables W1 (description of the sampling sites), W5 (miTAG related data) and W8 (a broad range of molecular concentrations, temperatures, and computational measures of diversity). The miTAG 16S abundance estimations were loaded into our R data frame as described in Methods 8. We first attempted to adjust for zeros in this dataset alongside our two samples using a multiplicate Bayesian approach^126^, but could not achieve convergence, since some Tara Oceans samples had zero counts for >50% of the taxa. To achieve convergence we had to remove taxa, a procedure we deemed undesirable and unnecessary as the observed adjustments were very small. We opted instead to simply add 1 to all entries in the count matrix.

Clustering was performed by first identifying the most abundant taxa for each sample, and then transforming these abundances to ranks. Then the Kendall T distance metric was used with Ward’s algorithm to construct two dimensional hierarchical clusters.

## Acknowledgements

We thank the Bellairs Research Institute of McGill University and the Centre for Resource Management and Environmental Studies (CERMES) at the University of the West Indies, Cavehill Campus, for their assistance with the logistics of experiments.

## Author Contributions

MH obtained the appropriate CITES permit for this work. RR, SM and VM assisted with the logistics of sample collection and transportation. MM and AR collected the samples taken at Maycocks and Bellairs respectively. SS and SK prepared samples for sequencing. HV and YV evaluated sampling, collected metadata and water chemistry. SS, VD and VB built the bioinformatic pipelines. SS, VD and MH performed data science analyses. DW provided guidance and financial support for sequencing. MH designed and supervised the project, contributing funding, analyses, and manuscript preparation. SS conducted the microscopy and measurement of physio-chemical variables in the study. This work was supported by grants awarded to MH from the NSERC Discovery and the Canadian Research Chairs programs.

## Data availability

The raw sequencing data is available at the National Center for Biotechnology Information (NCBI) Sequence Read Archive (SRA) with BioProject accession number PRJNA685579. Normalized sequencing data is available at the Joint Genome Institute (JGI) from the GOLD database with study ID Gs0136136. All code is available via GitHub at https://github.com/hallettmiket/barbadian-reef. This repository in turn provides access to all data and analyses at Zenodo.

This work was approved by the Coastal Zone Management Unit, of the Ministry of Maritime Affairs and the Blue Economy, of the Government of Barbados CZ01/9/9.

## Supplemental information: A metagenomic-based study of two sites from the Barbadian reef system

**Supplemental Figure 1.**
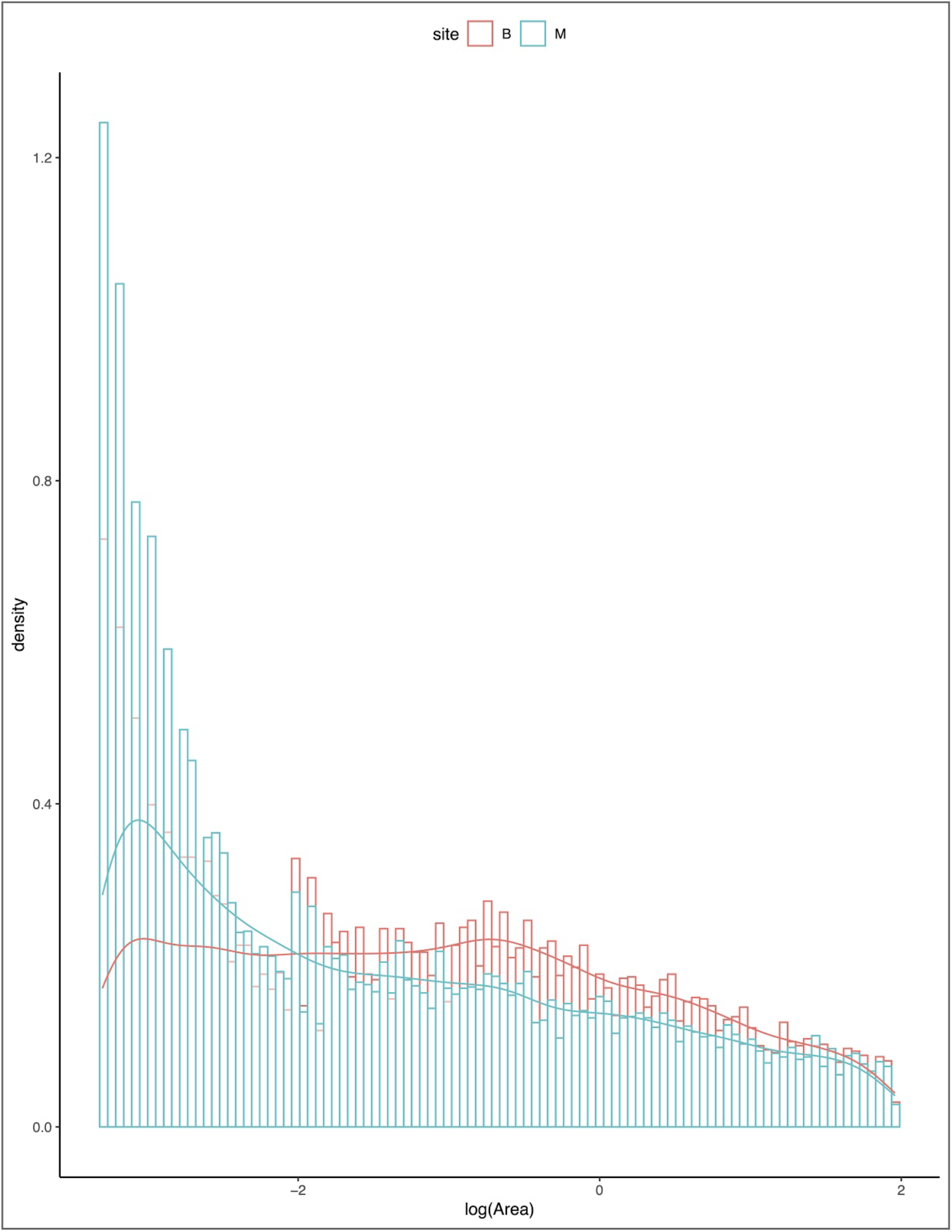
Distribution of the (log) area across ∼270K field of views (FOVs) from 60 images of reef water from Bellairs and Maycocks. We observe an enrichment of small organisms at Maycocks.

**Supplemental Figure 2.**
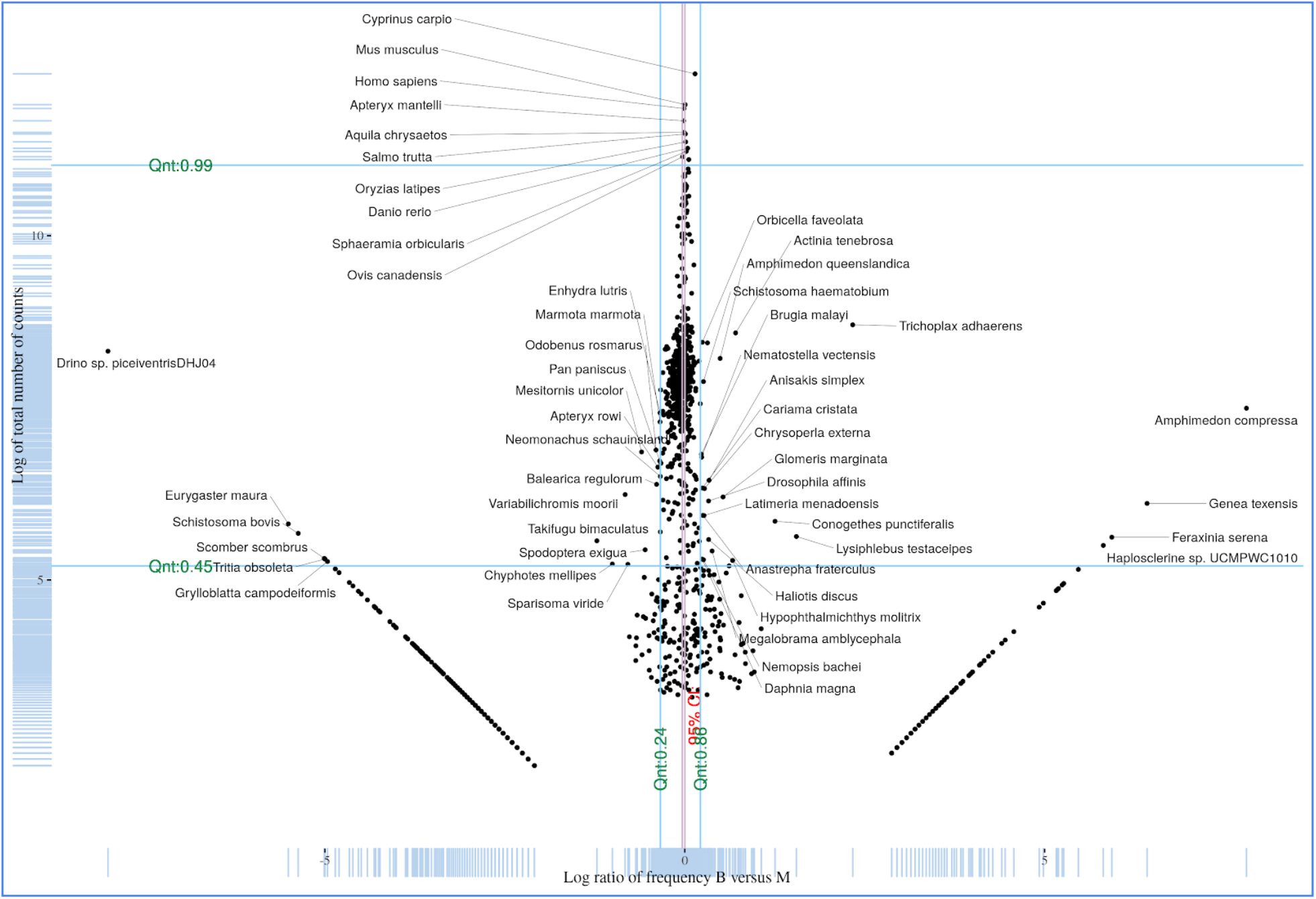
The distribution of Metazoan species. Here the x-axis corresponds to the log ratio of the fraction of reads mapped to a species at Bellairs (relative to the total number of reads mapped to Metazoa at Bellairs) versus the fraction of reads mapped to a species at Maycocks (relative to the total number of Metazoan reads at Maycocks). The y-axis is the log-sum of reads across both sites. Blue lines denote quantiles. The red lines denote a 95% bootstrap confidence interval for the mean of the distribution. The “v shape” in the lower left and right quadrants correspond to taxa identified uniquely at Maycocks and Bellairs respectively.

**Supplemental Figure 3.**
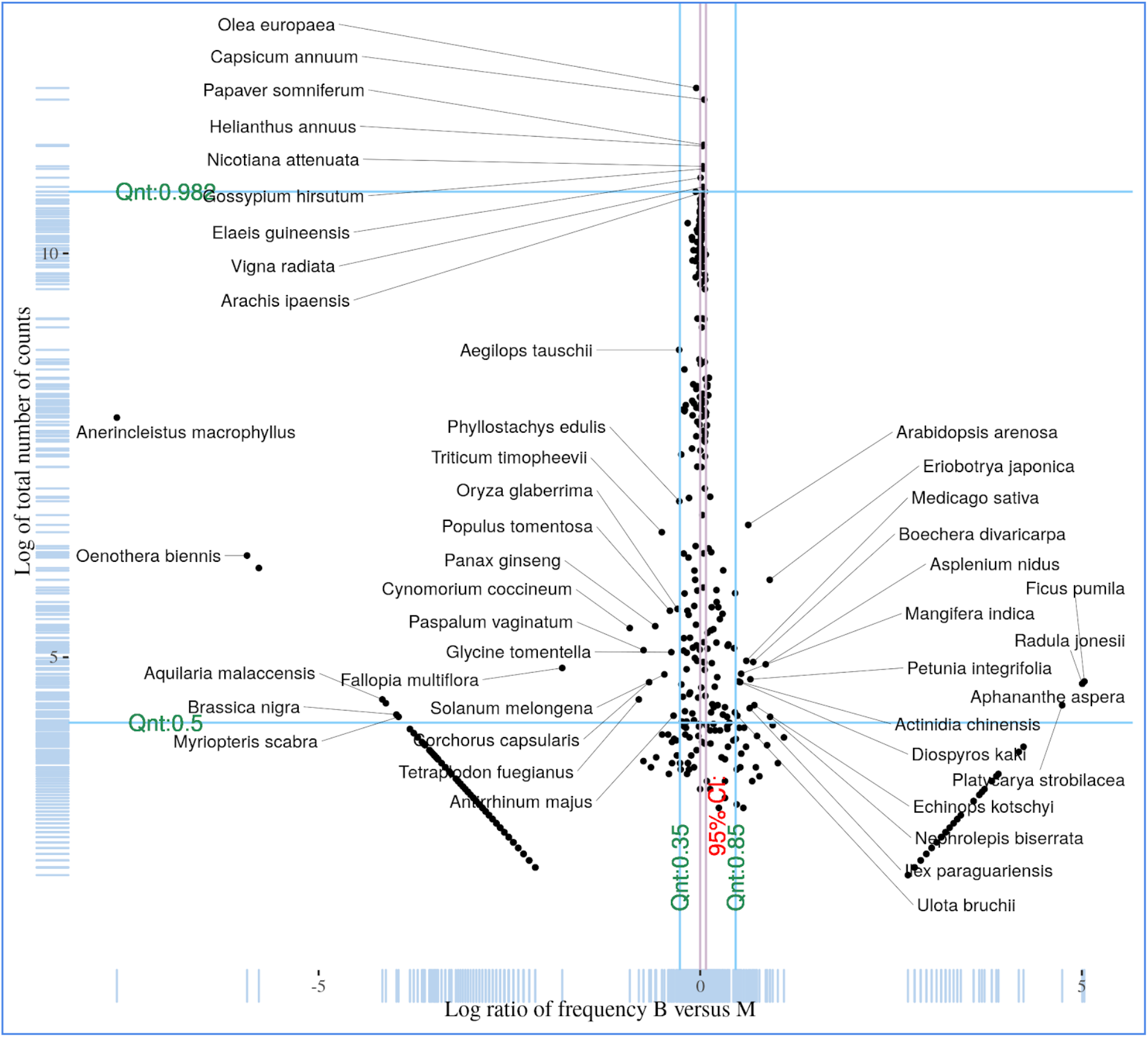
The distribution of Embryophyta species. Here the x-axis corresponds to the log ratio of the fraction of reads mapped to a species at Bellairs (relative to the total number of reads mapped to Metazoa at Bellairs) versus the fraction of reads mapped to a species at Maycocks (relative to the total number of Embryophyta reads at Maycocks). The y-axis is the log-sum of reads across both sites. Blue lines denote quantiles. The red lines denote a 95% confidence interval for the mean of the distribution.

**Supplemental Figure 4.**
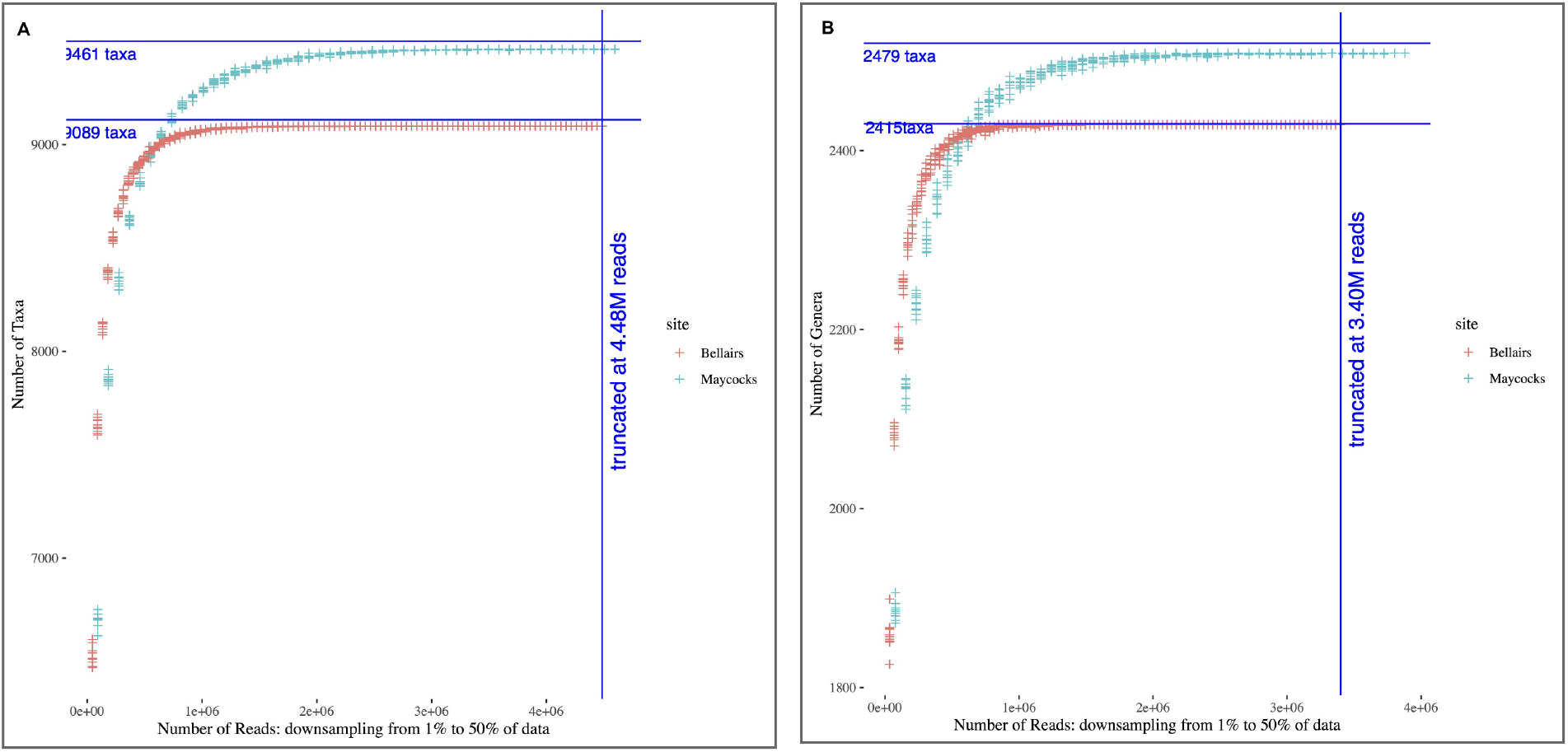
**A.** Downsampling of the paired-end reads highlighting the number of species identified. **B**. Downsampling of the paired end-reads highlighting the number of genera identified.

**Supplemental Figure 5.**
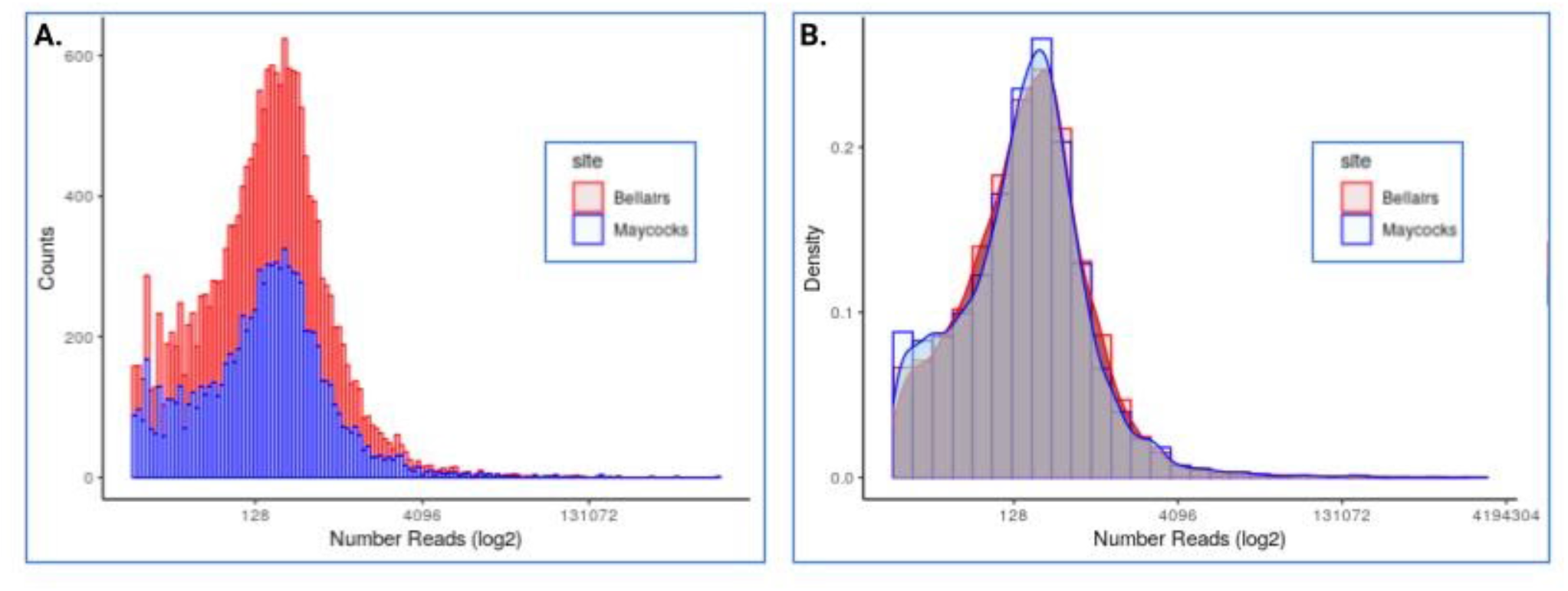
**A** The estimated number of species at both sites versus the log number of reads. **B** is the corresponding histogram of frequencies.

**Supplemental Figure 6.**
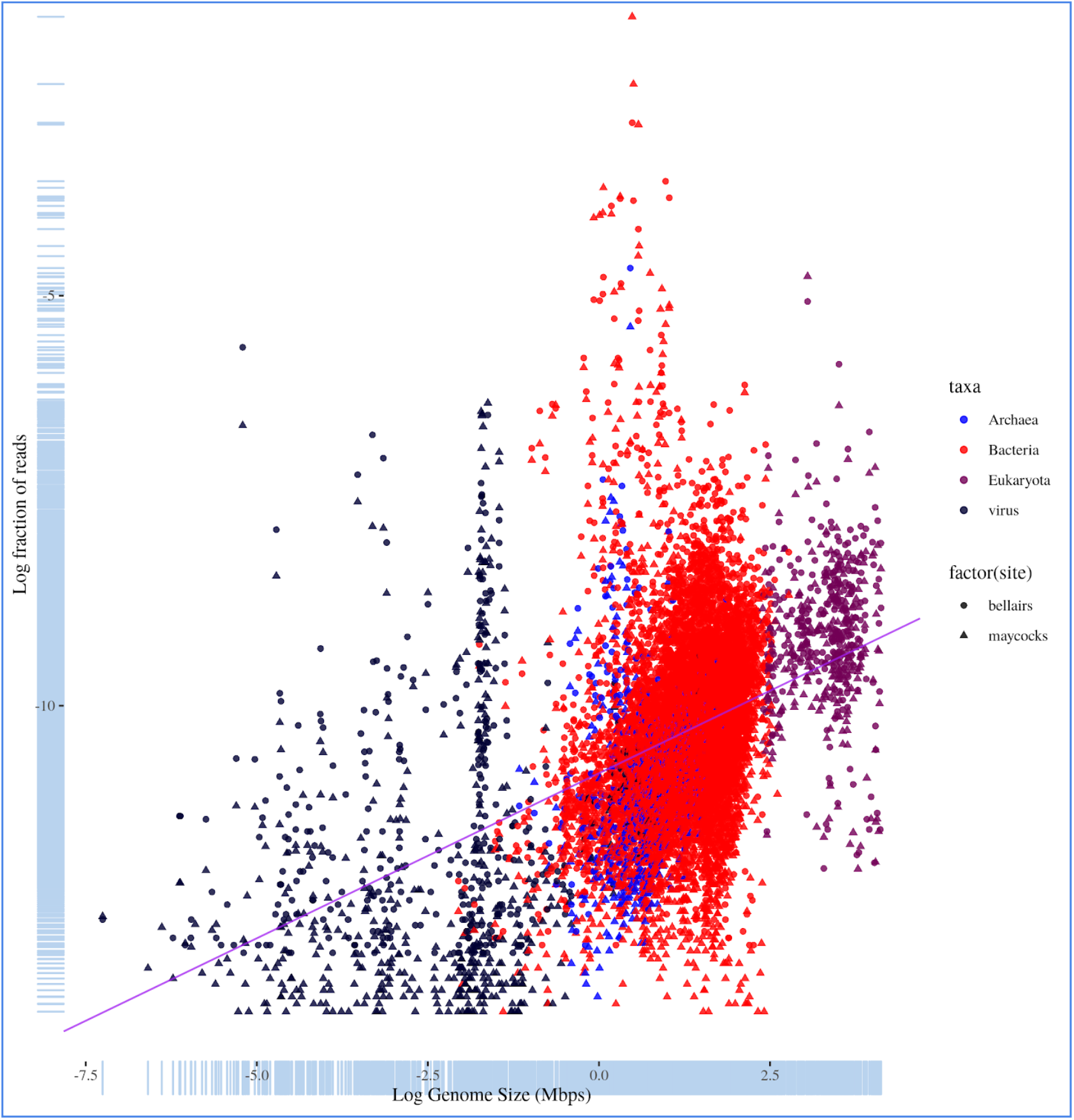
The relationship between genome size and number of reads mapped to the genome across all domains before correction.

**Supplemental Figure 7.**
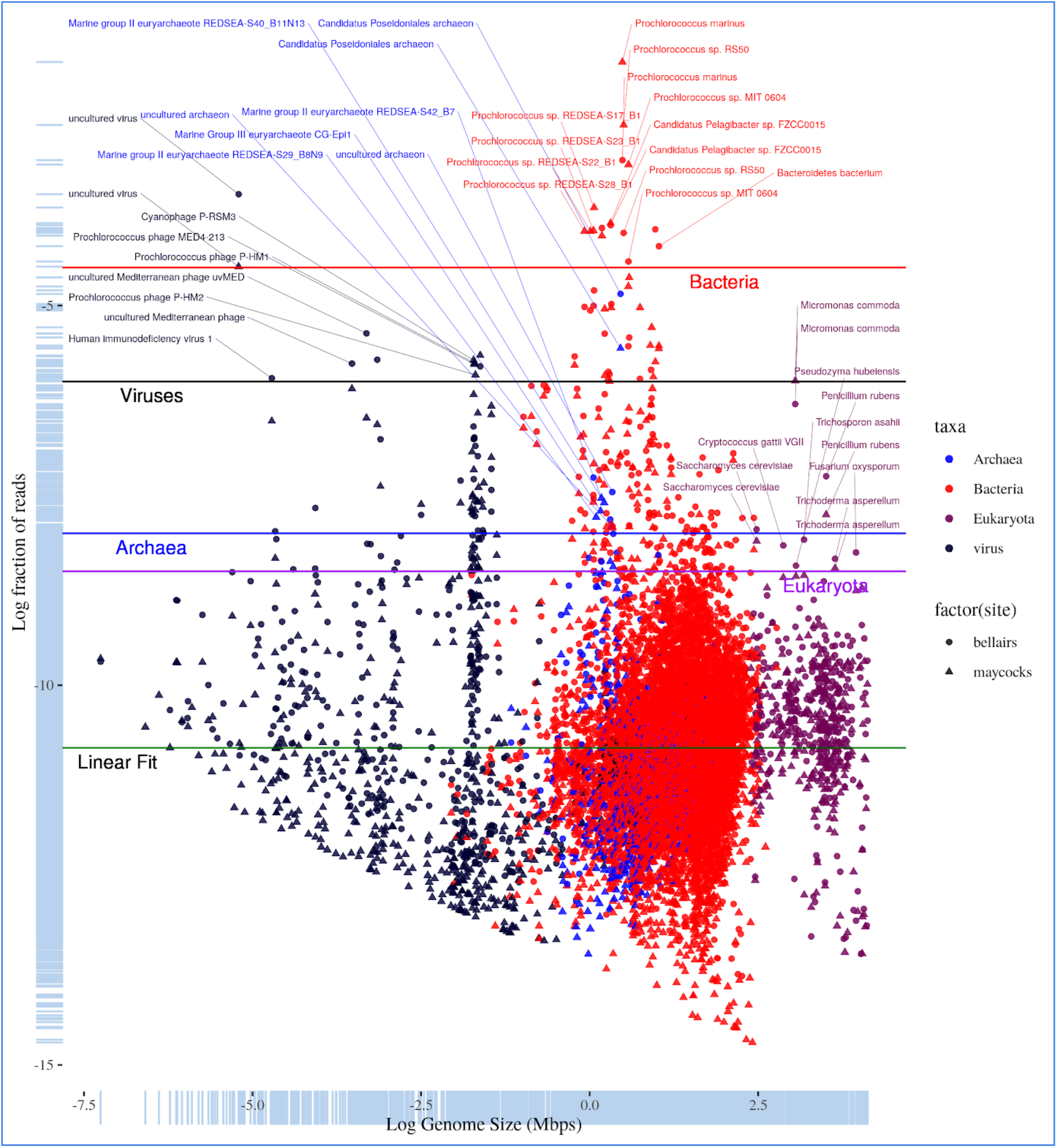
The relationship between genome size and number of reads mapped to the genome across all species across and all domains after adjusting by the slope obtained from a linear model across all species.

**Supplemental Figure 8.**
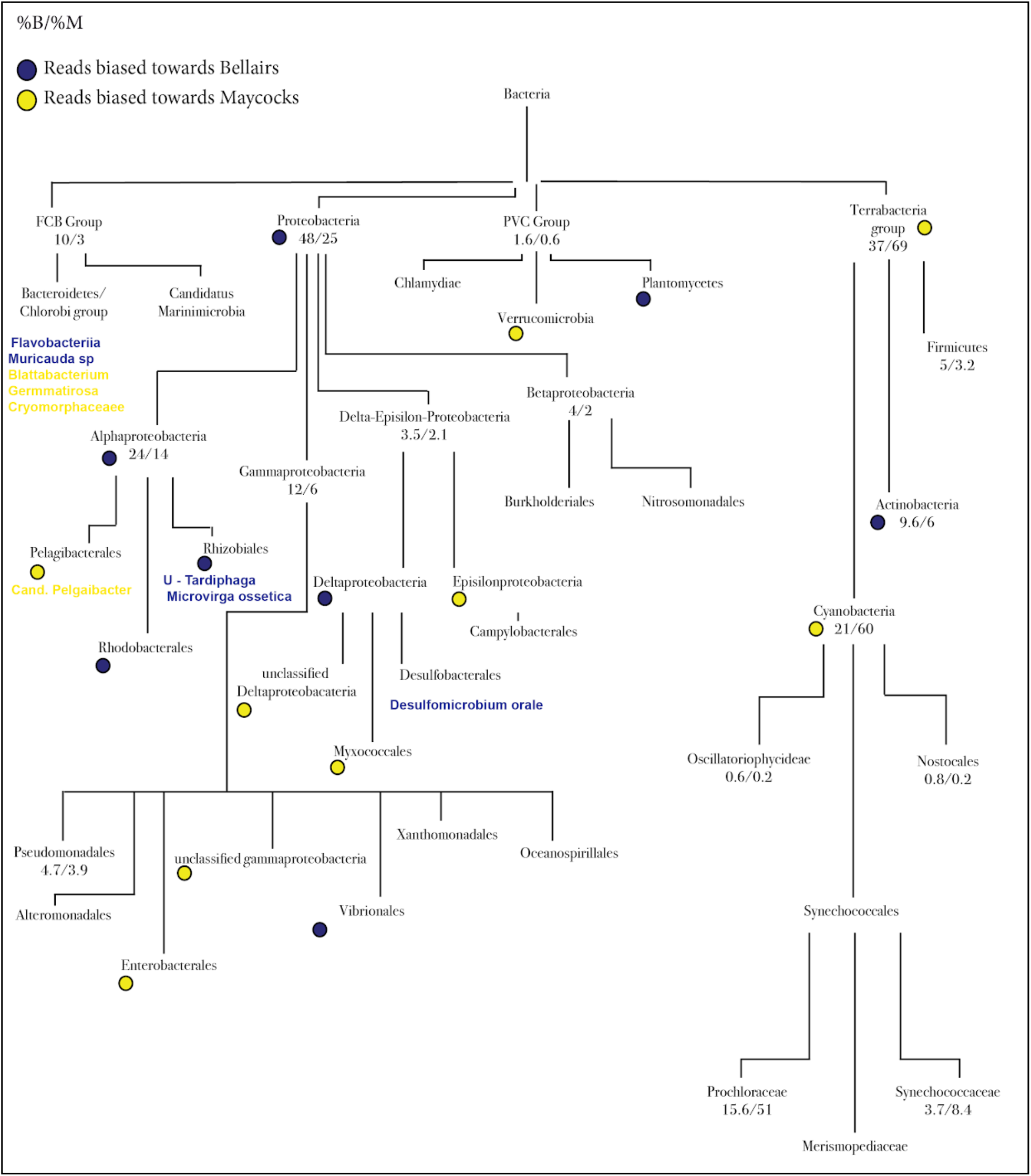
A schematic of the major Bacteria taxa identified at Bellairs and Maycocks. Here the numbers are the percentage of all bacterial reads mapped to the taxa (B/M). Blue and yellow circles indicate taxa where reads are significantly biased towards Bellairs and Maycocks respectively as determined via the KW test (p < 0.01).

**Supplemental Figure 9.**
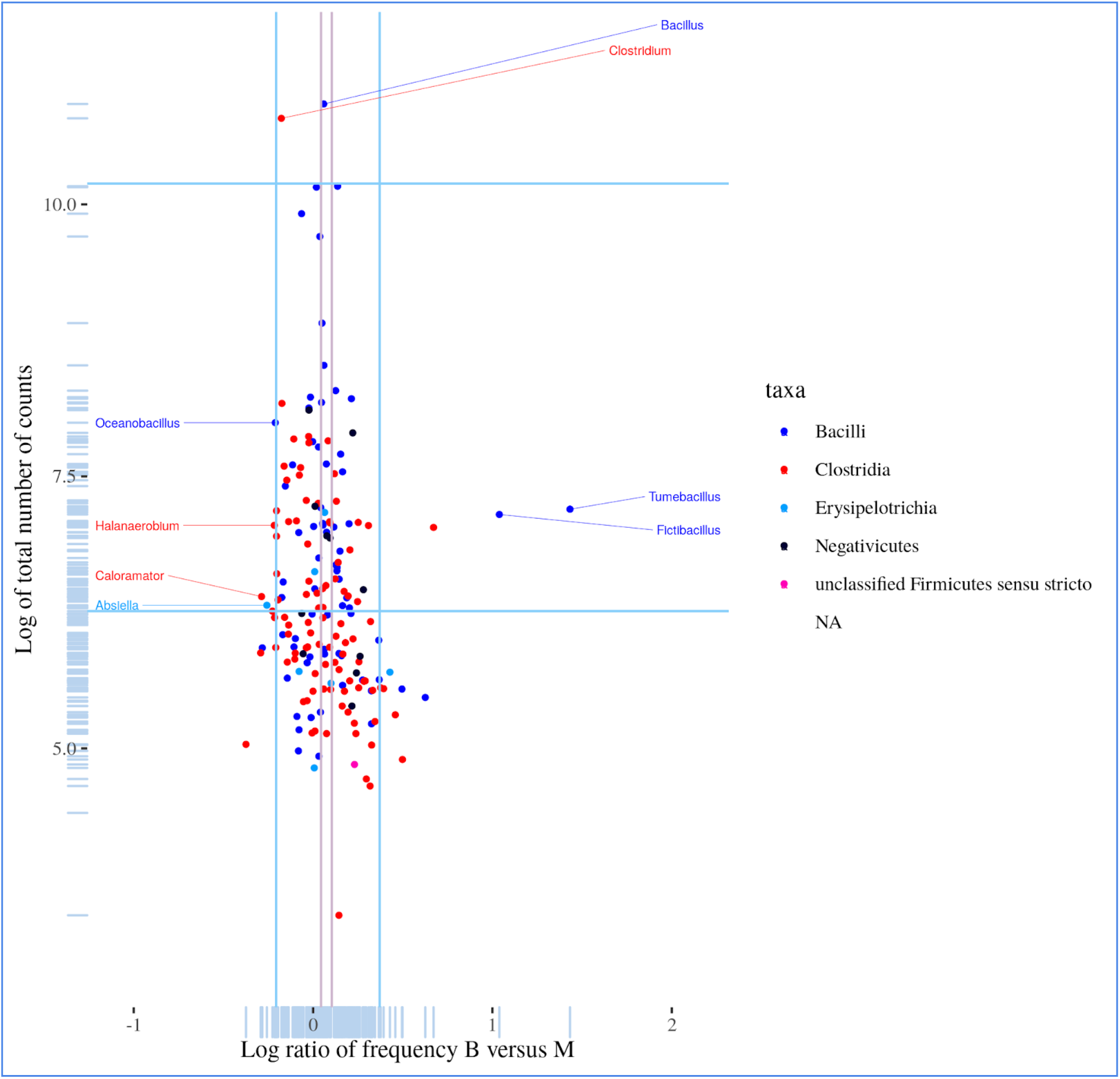
A log-log scatterplot at the genus level of Firmicutes.

**Supplemental Figure 10.**
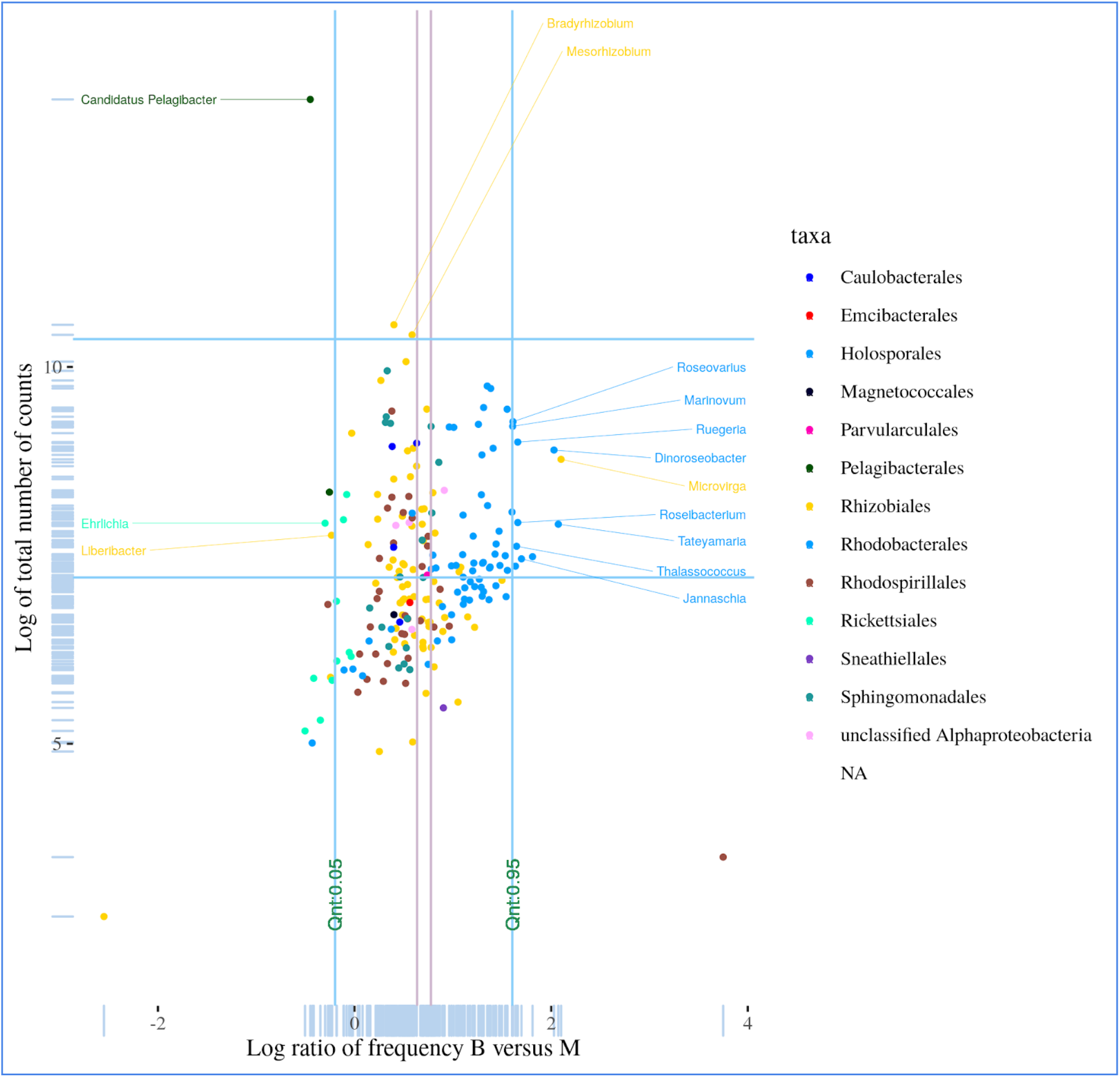
A log-log scatter plot of the log ratio of reads for all Alphaproteobacteria species versus the total number the total number of reads for the species across all of the bacterial domain but at the genus level.

**Supplemental Figure 11.**
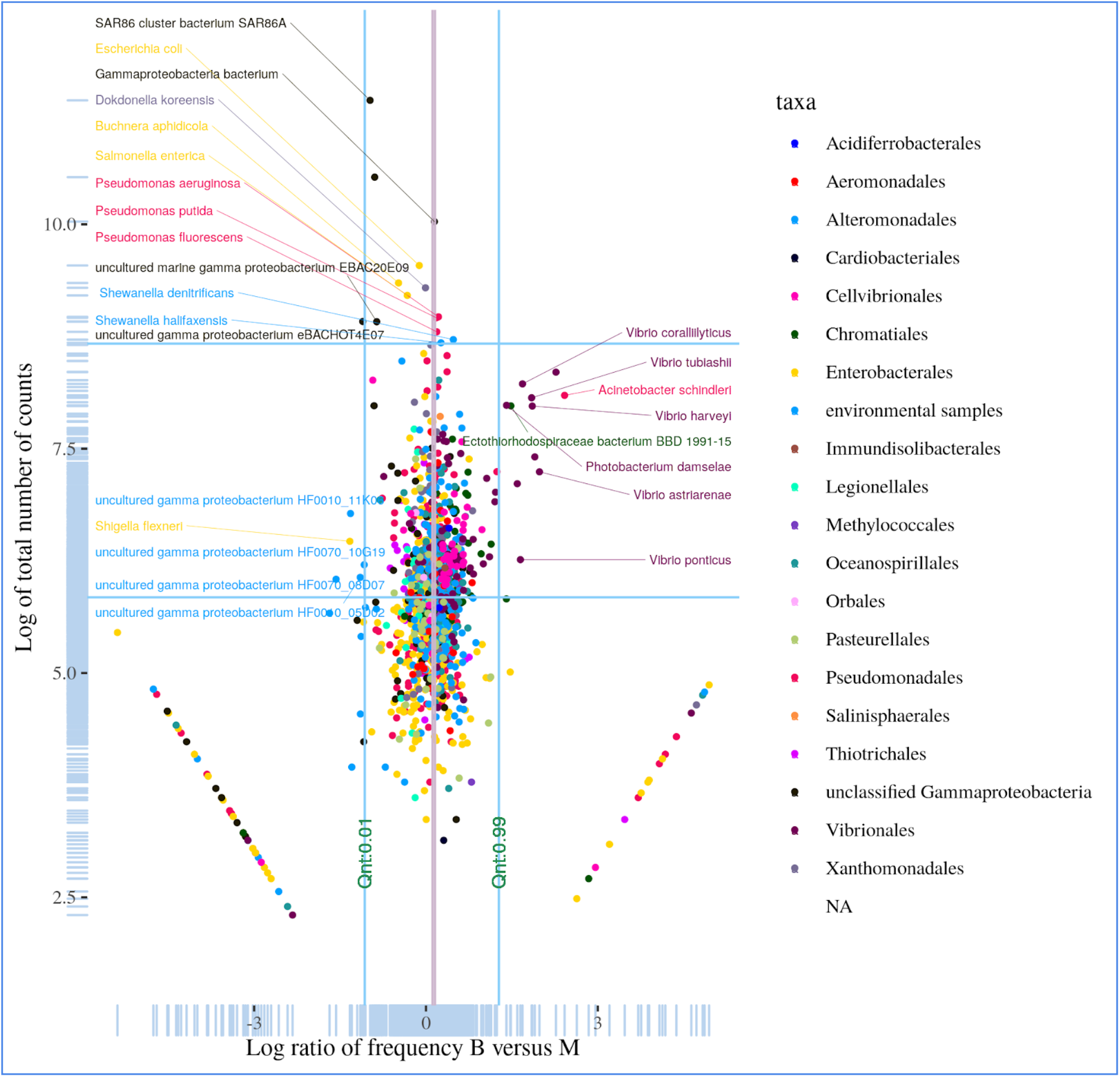
A log-log scatter plot of the log ratio of reads for all Gammaproteobacteria species versus the total number the total number of reads for the species.

**Supplemental Figure 12.**
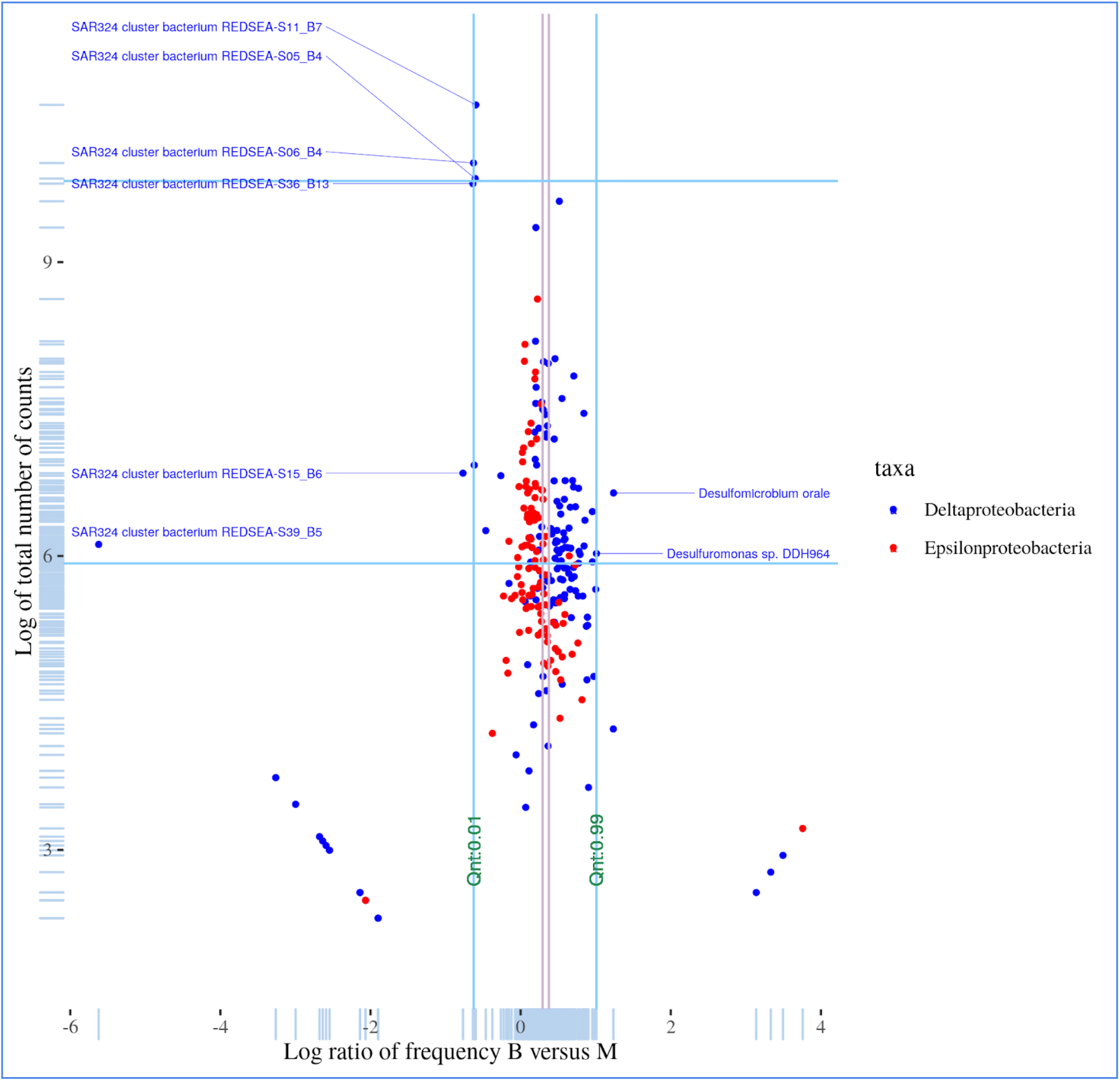
A log-log scatter plot of the log ratio of reads for all delta-epsilonproteobacterial species versus the total number the total number of reads for the species.

**Supplemental Figure 13.**
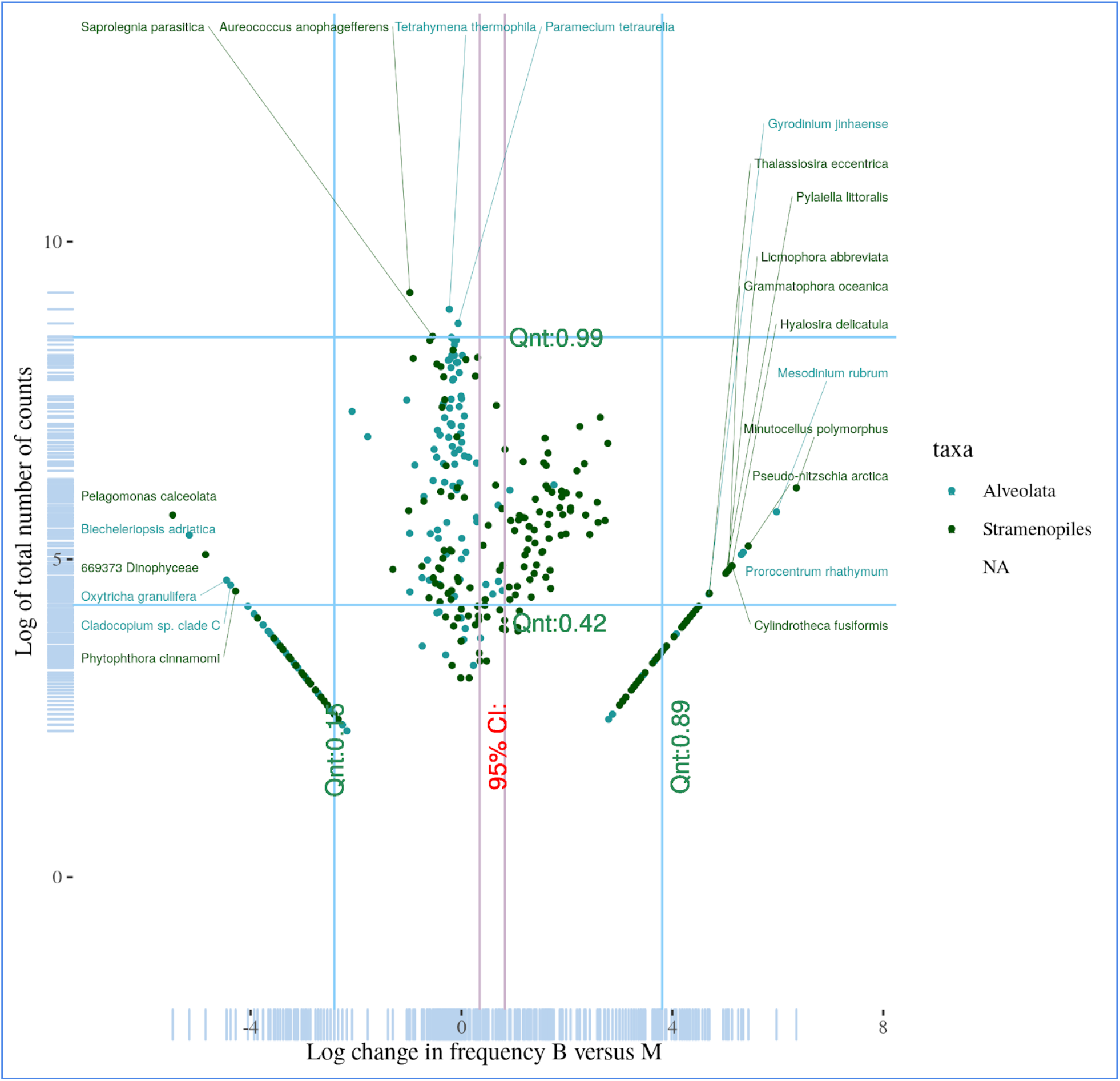
A log-log scatter plot of the log ratio of reads for all SAR species versus the total number the total number of reads for the species.

**Supplemental Figure 14.**
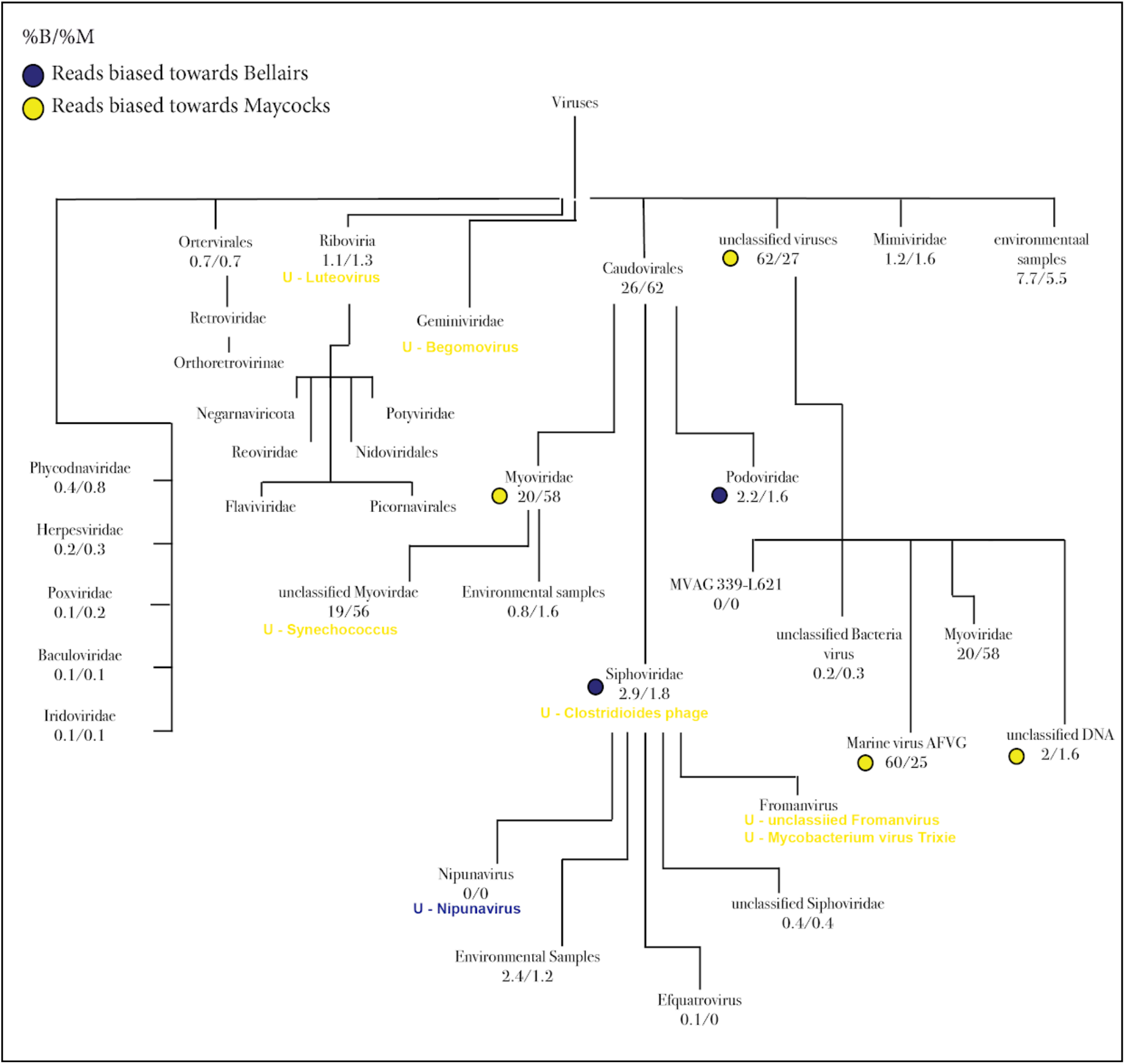
Schematic of the major taxa identified at Bellairs and Maycocks. Here the numbers are the percentage of all eukaryota reads mapped to the taxa (B/M). Blue and yellow circles indicate taxa where reads are significantly biased towards Bellairs and Maycocks respectively as determined via the KW test (p < 0.01). Species uniquely identified at a given site are denoted in blue for Bellairs and yellow for Maycocks.

**Supplemental Figure 15.**
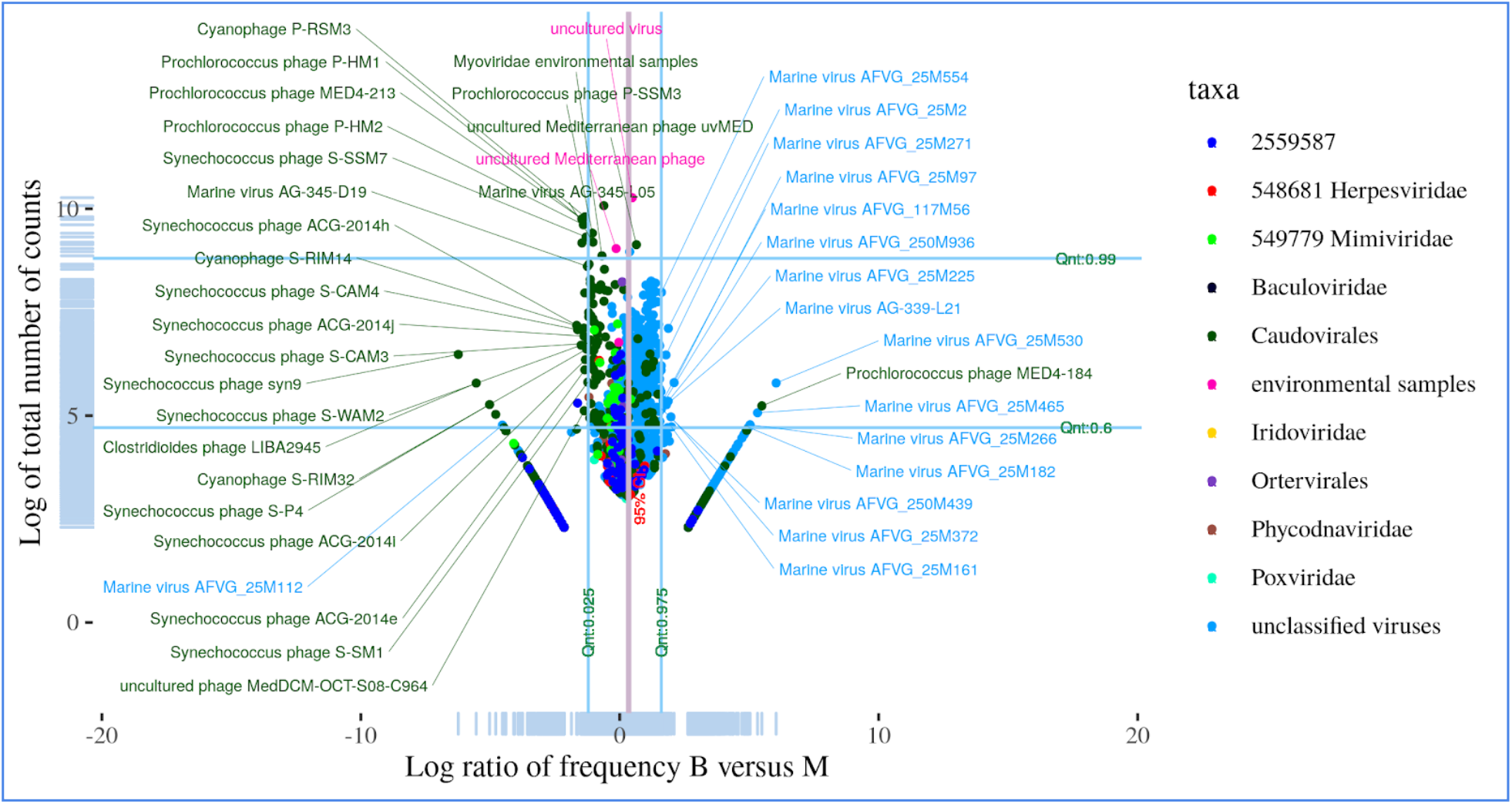
A log-log scatter plot of the log ratio of reads for all viruses versus the total number the total number of reads for the virus.

### Supplemental Information 1 - Profiling captured environmental DNA

Although our samples were filtered to remove large cells, we nevertheless observed reads mapping to non-microbial taxa such as Metazoa, Embryophyta, and multicellular fungi. The genomes of these organisms tend to be orders of magnitude larger in size than Bacteria and Archaebacteria. Therefore even a few such Eukaryotic cells escaping our filtering would provide a disproportionate amount of gDNA in our samples and garner a significant number of sequencing reads. Before removing these organisms from our analysis, we first asked what taxa were identified at the sites. Trace cells will likely not give accurate estimates of relative frequencies, but the existence of marine and related species likely indigenous to the reef system would provide supporting evidence towards the technical validity of our approach to detect marine species.

### Multicellular content: Metazoa are primarily marine-related and often indigenous to Barbados

An examination of Metazoa (**Supplemental Figure 2**) identified several species of fish at both sites including Taki fugu, Sparisoma viride, Latimeria menadoensis, and Megalobrama amblycephala. Although these taxa are indiginous to the Barbadian reef system, most of the remaining species identified did not have a clear link to Barbados, although they remained marine related. We hypothesize that very few reads mapped to very large genomes were not sufficient to reliably differentiate between lower levels in the tree of life. For example several other species of fish were identified including carp, salmon and atlantic mackerel. Several corals (eg Orbicella faveolata and Porites lobata), Cnidaria (Nematostella vectensis, Hydractinia symbiolongicarpus), sponges (Amphimedon queenslandica), tunicates (Halocynthia roretzi) shrimp (Macrobrachium nipponsense), a tunicate (Oikopleura doica), a sea snail (Haliotis discus), a water flea (Daphnia magna), the tortoise bug (Eurygaster master), several species of (marine) worms, and many basal free-living multicellular Eukaryota (eg Trichoplax adhaerens). Both sites had DNA from nematodes, snakes, insects (including several species of flies) and birds. Both sites had a significant number of reads mapped to human, green monkey (an old world primate common across the island), sheep, possum, and rock hyrax. The latter two species have close evolutionary relatives distributed across the island. Reads mapped to several non-indigenous marine mammals, especially at the Maycocks site including seal, walrus. Reads from all Metazoa were removed from our dataset, the relative frequencies of all taxa were recomputed.

### Multicellular content: Embryophyta correspond to marine, crop or ornamental plants common to the island

An examination of the multicellular Embryophyta (**Supplemental Figure 3**) identified several marine-related species including flowers (Papaver somniferum, Helianthus annuus) and trees. We stress again that the genomes of these taxa are very large but contribute just trace amounts of DNA to our samples. The number of reads is likely insufficient to identify the precise species or genera of tree. However, we did identify coastal salt-tolerant plants including Cynomorium coccineum and Paspalum vaginatum in the Maycocks sample, consistent with the observation that Maycocks has retained a wild grassy beach whereas the Bellairs beaches is adjacent to a research institute and hotels. The presence of Paspalum vaginatum on the island was reported recently and is likely associated with the development of golf courses^1^. DNA that may have originated from agricultural crops directly inland from Maycocks were detected including wheat/rich (Triticum timopheevil, Oryza glaberrima), ginseng (Panax ginseng), bamboo (Phyllostachys edulis), and jute (Corchorus capsularis). Several species were identified only at Bellairs including climbing fig (Ficus pumila), loquot (Eriobotrya Japonica), Platycarya tree (Platycarya strobilacea), tropical fern (Nephrolepis biserrata) and the muku tree (Aphanthe aspera). Other than the tropical fern (Nephrolepis biserrata) these species could not be directly identified as being present on shore at Bellairs. Some plant species that can be found on the shore of bellairs include West Indian mahogany (Swietenia mahagoni), flamboyant tree(Delonix regia), yellow-flamboyant (Peltophorum pterocarpum), Macarthur palm (Ptychosperma macarthurii) golden cane palm (Dypsis lutescens), manchineel tree (Hippomane mancinella), coconut tree (Cocos nucifera) and the whistling pine tree (Casuarina equisetifolia). Although not directly identified in our analysis some of these plant species that are present on the shore come from similar clades and orders of the species identified only at Bellairs. The species C. equisetifolia belongs to the Order Fagales of which P. strobilacea is also a member. While species S. mahagoni, D. regia, P. pterocarpum, H. mancinella belong to the Rosids clade of which E. japonica and A. aspera are members of. We note that sequencing at the Bellairs site produced far fewer overall reads that Maycocks. Therefore, even for a rare species, it is unlikely to be detected at Bellairs but not at Maycocks. After removing these taxa, we also removed reads mapped to several multicellular fungi and algae, in addition to plasmids and artificial sequences.

### Supplemental Information 2: Richness, diversity and handling zeros

Biodiversity can be measured in many different ways including taxonomic diversity (the presence of different species), phylogenetic diversity (the presence of different evolutionary lineages), or functional diversity (the variety of growth forms and resource use strategies)^2^. *Abundance* refers to the fraction of each species at a site, and since we are in CoDa setting, this is inherently a relative abundance. Diversity is a measure of the distribution of these abundances. Throughout this manuscript we use the Shannon index as our measure of diversity^3^ with a permutation based approach to estimate p-values

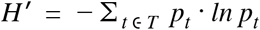

where *T* is the vector of relative abundance for all taxa and *p*_*t*_ is the probability of taxa *t*. Briefly, there is an increase in the Shannon index (entropy) as the distribution of abundances approaches a “flat” uniform distribution, and decreased entropy as the frequency of one (or a few species) approaches one.

*Richness* is defined as the number of species within a specified clade at a site. We asked if there was a difference in richness between Bellairs and Maycocks. However, we must first adjust for the fact that Maycocks and Bellairs had different levels of sequencing coverage. More specifically, budget and technical limitations imply that the number of draws made by the sequencer from the urn is finite although large. The sequencing coverage (total number of reads) may not be sufficiently large to identify with high probability rare species in the sample. For instance, a species whose DNA contributes only 1 read to an urn with 10M reads is unlikely to be identified, if the sequencing coverage is only 1M. In our data, we obtained ∼4.5M and ∼9.2M mappable paired-end reads from Bellairs and Maycocks respectively. Therefore, we expect to identify more taxa in Maycocks than in Bellairs due to this reason alone. In other words, species richness increases with sample size, and differences in richness may be due to differences in sample size.

To address this, we first downsampled from the Illumina paired-end reads from 1-50%, and counted the number of identified species and genera (**Supplemental Figure 4**). After only ∼3M reads at Maycocks and ∼1.5M reads at Bellairs (< of total in both cases), all taxa have been identified. Maycocks however converges 272 more species (64 more genera) than Bellairs.

Rarefaction provides a second approach to exploring this issue. A statistical correction is computed that estimates the number of taxa we would have observed at the sites if we had sequenced Maycocks to the same coverage as Bellairs. To rarefy samples from N to n total reads, we used the rarefy function in the vegan package^4^. In particular, we extrapolated the number of species at each site individually and also with pooled data using the prestondistr function to estimate the log-normal fit, and the veilespec function to estimate the integral fitted with `prestondistr`. Consistent with your downsampling approach, this statistic suggested that no change in the number of identified species. This suggests our sequencing coverage is sufficient for all but the most extremely rare organisms.

In ecological communities including marine, most species are rare^5^. Preston argued that this implies that richness would follow a truncated log normal distribution (Preston 1948). This is true for the relative frequencies obtained for our data as depicted in **Supplemental Figure 5**. This allows us to estimate the theoretical richness at both sites, the so-called Preston veil^4,6^. Specifically, by integrating the fitted log-normal, the Preston veil speculates how long the right tail is if we had infinite sequencing data. However, consistent with the analysis above, the Preston veil did not predict any new species would have been identified (it predicted 0.28 more species above the 9089 species observed). The results for Maycocks were equally insignificant. If there is a large difference in the sequencing coverage between sites (as is our case), the presence zeros in the shallower site can have non-intuitive effects on analyses especially with respect to distance measures and clustering. Moreover, CoDa analysis (such as ours here) often relies on log-ratios of the form 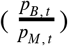 where *p*_*B,t*_ and *p*_*M,t*_ are the estimates of the frequency of taxa *t* at Bellairs and Maycocks respectively. This is referred to as the zero handling problem^7^. We applied a Bayesian-multiplicative replacement strategy that adjusts the count matrix (for all taxa at both sites) in a manner that preserves the ratios between the non-zero components^8^. However, when external datasets (eg Tara Oceans) were included in this analysis with lower sequencing coverage, they had > 50% zeros. We were not able to reach convergence unless we removed taxa. After removing taxa that had zero counts in more than 50% of the samples, we reached convergence but the adjustment had little effect on the data.

### Supplemental Information 3: Correcting for genome size

We asked if there was a correlation between genome size and number of reads aligned to each species across Archaea, Bacteria, Eukaryota and Viruses. The log-log scatterplots of **Supplemental Figure 6** depicts these relationships across all species. Visual inspection suggests a correlation between log genome size and log read count as expected. To adjust for this effect, we fit a simple linear model of the form

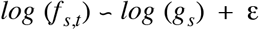

where f_s,t_ is the fraction of all reads mapped to taxon t at site s, and g_s_ is the genome size (Mbp) for species s, and E is a normally distributed random variable. An implicit assumption in this simple model is that the vast majority of taxa have approximately the same fraction of read counts f_s,t_. The parameters of the fit were then used to correct the observed read counts. Given the compositional nature of our data, any investigations in this manuscript that seek to compare two taxa *within* the same site must first adjust read counts using the linear model.

The corrected relative abundance estimates of all species are depicted in **Supplemental Figure 7**. The adjustments highlight a high abundance of the bacteria Candidatus Pelagibacter and Prochlorococcus. Several Archaea including the Marine Group II/III and Cand. Poseidoniales appear on magnitude below, followed by viruses that inflect Prochlorococcus and lastly one the Eukaryota Micromonas commoda appear.

### Supplemental Information 4: Viruses

Marine viruses affect microbial populations by releasing carbon and nutrients into the ecosystem through lysis, by complexing nutrients such as iron, through reprogramming of host metabolism and horizontal gene transfer, and via the formation of relationships including for example cyanobacteria-cyanophage relationships which affect CO2 fixation ^9-13^. Metagenomic analysis of marine viruses has been investigated including in the context of the Tara Oceans Project ^11^. Brum and colleagues isolated organisms below 0.22 µm and built optimized sequencing and bioinformatics platforms specific for the analysis of viromes. Our investigation here is limited, since we selected for organisms between 0.22 µm and 3 µm. Nevertheless, 5.8% (B) and 3.6% (M) of all reads mapped to viruses (**Supplemental Figure 14**). Brum and colleagues also built specialized analytic pipelines; we however use the same pipeline for our preliminary investigations here.

### Viruses: Bellairs is highly enriched for uncharacterized phages

A large fraction of all viral reads were mapped to virus genomes reported first in a study that developed an assembly-free single molecule nanopore sequencing approach for viruses from environmental samples obtained close to Hawaii^14^. These so-called assembly-free virus genomes (AVGVs) show a clear preference for Bellairs (KW, p << 0.01). Moreover, the original virus-enriched samples sequenced by Beaulaurier were collected at 25, 117, or 250 meters with n=565, 93, 1023 respectively. The samples from Beaulaurier et al. were filtered to remove all organisms > 0.22 µm. Of the 14 Marine virus AFVG in the 97.5% percentile across all viruses in our data (**Supplemental Figure 15**), 11 were harvested at 25 meters. This depth is closest to our samples harvested just below the surface. In Beaulaurier et al, they estimate that 26% of the AFVGs from 25m correspond to cyanophages, 13.3% to SAR11 phages, 12% to SAR116 phages, and 3% to Vibrio phages.

### Viruses: Cyanophages are highly enriched at Maycocks, a site highly enriched for cyanobacteria

Consistent with the strong preference for Cyanobacteria (incl. Prochlorococcus and Synechococcus) at Maycocks (26% B vs 62% M), there is a comparably strong preference for cyanophages at Maycocks (KW and Dunn’s test for Caudovirales, both p << 0.01, **Supplemental Figure 15**). Consistent with previous findings^15^, Myo-, Sipho- and Podo-viruses are found in our data; these are well established phages for Prochlorococcus and/or Synechococcus.

### Viruses: Bellairs is enriched for phages of Pelagibacter

Although the small heterotrophic Pelagibacter, a member of the ubiquitous SAR11 clade, is highly enriched at Maycocks, Podovirus phages of Pelagibacter are systematically shifted towards Bellairs (KW, Dunn’s test, p<<0.01).

### Viruses: Bellairs is enriched for plant, animal and algae viruses

Several additional plant, animal and algae viruses were enriched at Bellairs including Negarnaviruses (Canine morbillivirus, Salmon isavirus), Potyviruses (Turnip mosaic virus, Sugarcane mosaic virus), Phycodnaviruses and Iridoviruses.

### Viruses: Comparison with the Tara Oceans data

We compared the Shannon index of the Myoviridae, Podoviridae and Giruses against the analogous values reported for the Tara Oceans data^16^. At 11 degrees latitude, Tara Oceans data established the entropy at 3+/- 0.2 for Myoviridae, whereas we report a slightly higher entropy at 3.68 (B) and 3.63 (M). For Podoviridae, Tara Oceans reports 5 +/- 0.1, whereas we have a much smaller entropy at 2.94 (B and M). The giant viruses (giruses) have an entropy of 5.5 +/- 0.1 in the Tara Oceans data. As a working definition for viruses, we included any virus specus to the nucleocytoviricota clade. In our data the Shannon index is much lower at 3.19 (B) and 3.2 (M).

